# Germline removal reprograms somatic genome maintenance towards faster, energy-efficient, high-fidelity DNA defence and repair

**DOI:** 10.64898/2026.09.09.750329

**Authors:** Jean-Charles de Coriolis, Ghazal Alavioon, Wilfried Haerty, Emmanouil Tsakoumis, Monika Schmitz, Alice M. Godden, Raheleh Rahbari, Alexei A. Maklakov, Simone Immler

## Abstract

Germ cells maintain their genomes with greater fidelity than somatic cells, yet how germline status systemically shapes somatic genome protection in vertebrates remains poorly understood. Combining germline ablation with multi-organ transcriptomics in zebrafish (*Danio rerio*), we dissected germline control of somatic DNA repair and genotoxic stress responses. Using CRISPR/Cas9-mediated knockout of the germline determinant gene *dnd-1*, we compared germline-free (GLF) males and germline-carrying (GLC) siblings across five somatic organs at baseline and following sub-lethal γ-irradiation, with sampling at 3 h and 24 h post-irradiation.

Germline ablation profoundly reorganised baseline somatic transcription, favouring shorter, exon-rich genes with functions in chromatin integrity, cell-cycle control, and high-fidelity DNA repair, and attenuating DREAM complex repression of canonical repair targets. After irradiation, GLF fish mounted a faster response that rapidly engaged homologous recombination, checkpoint control, and proteostasis, whereas GLC fish remained biased toward biosynthetic and translational programmes. Hedgehog signalling acted as a transient regulatory node coordinating the strain-specific DNA damage response, and a repeat-rich region of chromosome 4 emerged as a hotspot of coordinated gene–transposable element (TE) activity. GLF soma showed reduced early TE expression after irradiation, with LTR retrotransposons showing the strongest concordance with nearby differentially expressed genes. Our findings identify germline status as a systems-level switch that tunes somatic genome maintenance and TE dynamics across multiple organs and provide a molecular framework for understanding how the germline regulates somatic ageing in vertebrates.

## Introduction

Cellular genomes experience continuous exogenous and endogenous stress, producing DNA lesions that are countered by dedicated repair pathways to preserve genome integrity^1^.

Nevertheless, repair is imperfect, and residual errors generate mutations that undermine genome stability and cellular function. Across the animal kingdom, germ cells deploy more accurate repair mechanisms than somatic cells, which rely more heavily on error-prone pathways; consequently, somatic mutation rates exceed germline rates. Diminished repair capacity in somatic tissues drives age-associated accumulation of mutations^2^ and genome instability, and DNA repair capacity itself declines with age^3^. Beyond fuelling oncogenesis through mutation accumulation, DNA damage in somatic cells can stall transcription and replication, broadly disrupting cellular processes. DNA damage thus likely contributes to organismal ageing through perturbed signalling, loss of proteostasis, and impaired mitochondrial function.

The germline influences somatic maintenance, including somatic DNA repair, and germline ablation can increase stress resistance and longevity^4^. In response to genotoxic stress, the germline can enhance somatic protection^5^. The “disposable soma” hypothesis posits a resource-allocation trade-off between reproduction and somatic maintenance^6,7^: organisms may favour less costly but more error-prone repair in soma to conserve resources for germline fidelity^8^. Consistent with this view, germline ablation accelerates somatic repair after genotoxic stress in zebrafish^9^.

However, other studies indicate that the negative association between germline and somatic maintenance can be uncoupled^8,10^. Removal of the somatic gonad eliminates reproduction but does not extend lifespan in *Caenorhabditis elegans*, whereas germline removal does, implicating the somatic gonad in endocrine signalling that modulates lifespan. Similarly, in killifish *Nothobranchius furzeri*, germline depletion enhances somatic maintenance after genotoxic stress, whereas arresting germline differentiation does not^4^, suggesting that germline-derived signals, rather than reproduction *per se*, govern somatic repair. In both dioecious nematodes *C. remanei*^11^ and killifish *N. furzeri*^4,12^, germline removal increases male lifespan but not female lifespan, further arguing against a universal resource-allocation explanation.

These observations point to cellular signalling pathways that tune somatic DNA repair in response to environmental context. When environmental stress compromises the germline or precludes immediate reproduction, germline signals may promote heightened somatic genome maintenance to sustain survival^5^. More broadly, when the environment suppresses fertility, selection for survival under diverse conditions may intensify^13^. Resource allocation and endocrine signalling hypotheses are not mutually exclusive, and the costs of germline maintenance can be sex- and environment-specific. The activity of transposable elements (TEs) - a major source of double-strand breaks - is markedly stronger in the germline than in soma^14^. Comparing TE activity across somatic organs and germline under varying environments therefore provides a window into somatic maintenance via germline signalling.

Here, we examined germline regulation of somatic genome maintenance and repair across five somatic organs in male zebrafish. We compared the impact of germline ablation on somatic maintenance at baseline and after non-lethal γ-irradiation, a genotoxic stress that elicits robust responses in both germline and soma. We profiled transcriptomes and TE expression in brain, intestine, kidney, spleen, and caudal fin and, when available, testes, comparing germline-free males (GLF; homozygous *dnd-1* knockout) with germline-carrying full-sib controls (GLC; homozygous wild type). Tissues were sampled at 3 h and 24 h post-irradiation (hpir) to capture temporal dynamics of the response. We found that germline ablation broadly upregulates somatic genome protection and repair under standard and stressful conditions, accelerates the response to genotoxic stress, and alters TE expression in the soma.

## Results

### Germline ablation reorganises baseline somatic transcription

Across five somatic organs, GLF and GLC males exhibited extensive baseline transcriptional divergence. We detected 6,855 differentially expressed genes (DEGs), of which 4,498 were upregulated in GLC and 2,357 in GLF (Supplementary Tables S1–S2). More than 270 Gene Ontology (GO) terms were shared between strains, largely related to transcriptional regulation, genome maintenance, and protein synthesis, but the majority were upregulated in GLC soma (Wilcoxon *V* = 1,701.5, *p* < 0.0001; Supplementary Figure S3). Semantic similarity analysis revealed low overall overlap of gene sets underlying shared GO terms, with greater similarity for double-strand break repair and ribosome biogenesis and lower similarity for microtubule organisation, DNA replication, and chromatin remodeling (Supplementary Figures S2, S3, S5).

Functionally, GLF-upregulated genes were enriched for cell-cycle progression and chromatin-linked DNA integrity (e.g., telomere maintenance, DNA packaging), whereas GLC-upregulated genes emphasised metabolic activity and adaptive stress responses (e.g., mitochondrial gene expression, ubiquitin-mediated proteolysis; Supplementary Figure S3). Organ-specific analyses recapitulated these trends, with heightened genome maintenance signatures in brain and kidney, immune activation in intestine and spleen, and structural/checkpoint control in caudal fin (Supplementary Figure S3). The top DEGs included *armc9*, involved in ciliogenesis and cell–cell communication, and *nlrc11*, involved in antiviral immunity, both upregulated in GLF soma; whereas *mhc2dab* and *sppl2*, both immune and differentiation genes, were upregulated in GLC soma (Supplementary Figure S2; Supplementary Tables S3–S5).

Global properties of DEGs differed significantly between strains (Figure 2), as indicated by a significant interaction between contrast and strain for gene expression (χ*²* = 2,252, df = 17, *p* < 0.0001), gene length (χ*²* = 1,538, df = 17, *p* < 0.0001), and exonic density (χ*²* = 301,334, df = 17, *p* < 0.0001). In GLC, DEGs were longer, had lower exonic density, and lower expression relative to the transcriptome average (Supplementary Tables S6–S7), whereas in GLF, DEGs were shorter and exon-rich, consistent with reduced transcriptional expenditure and increased splicing efficiency^15–17^.

**Figure 1.**
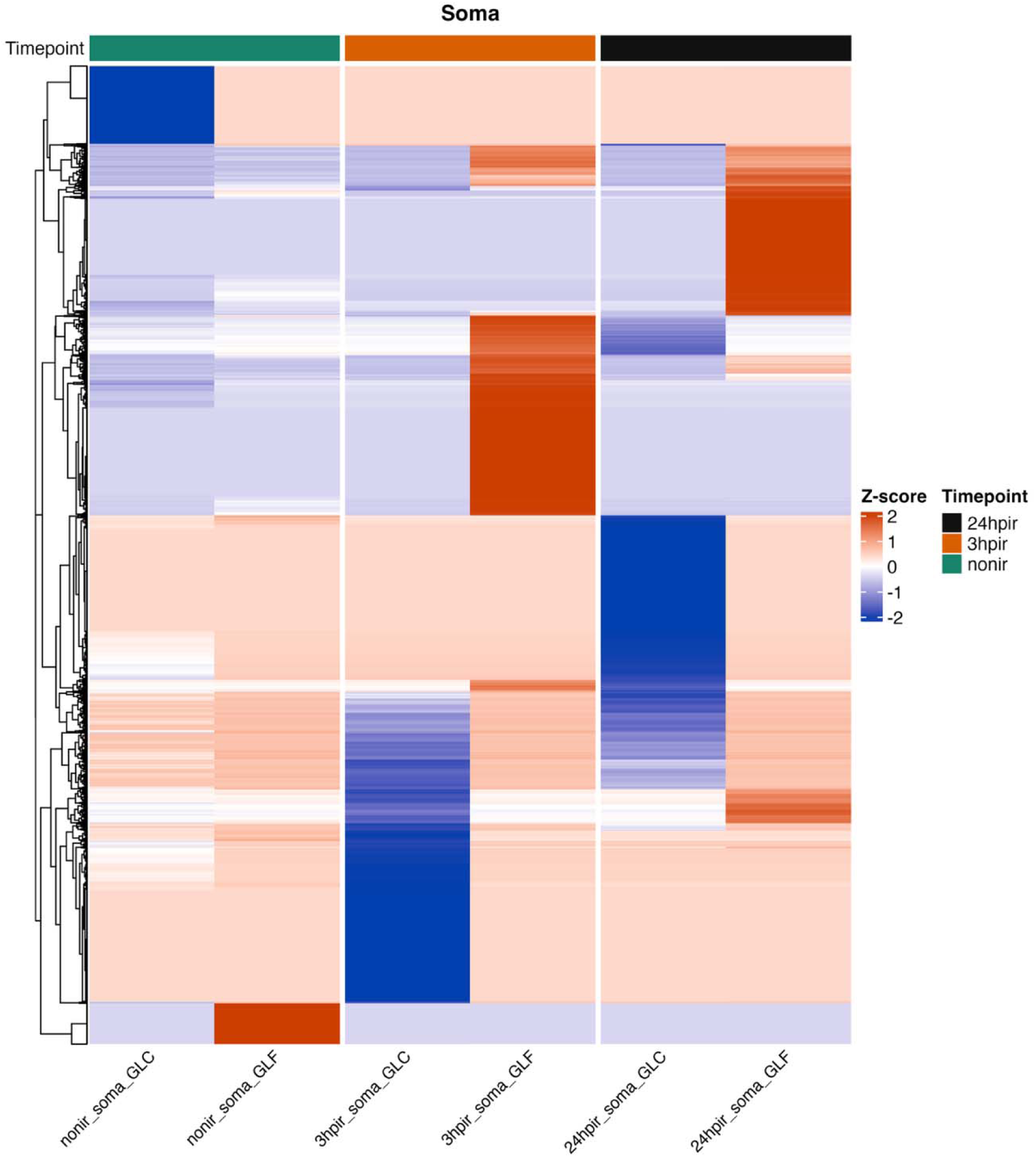
Global transcription landscape of somatic tissues reveals strain- and irradiation-dependent gene expression patterns. Gene expression values (logCPM) were normalised per gene across somatic tissues of GLC and GLF fish to compute Z-scores representing relative expression levels. Positive *Z*-scores (orange shading) indicate expression above the gene’s mean, negative *Z*-scores (blue shading) indicate below-mean expression. Rows correspond to individual genes, columns represent samples grouped by strain and timepoint post irradiation.

**Figure 2.**
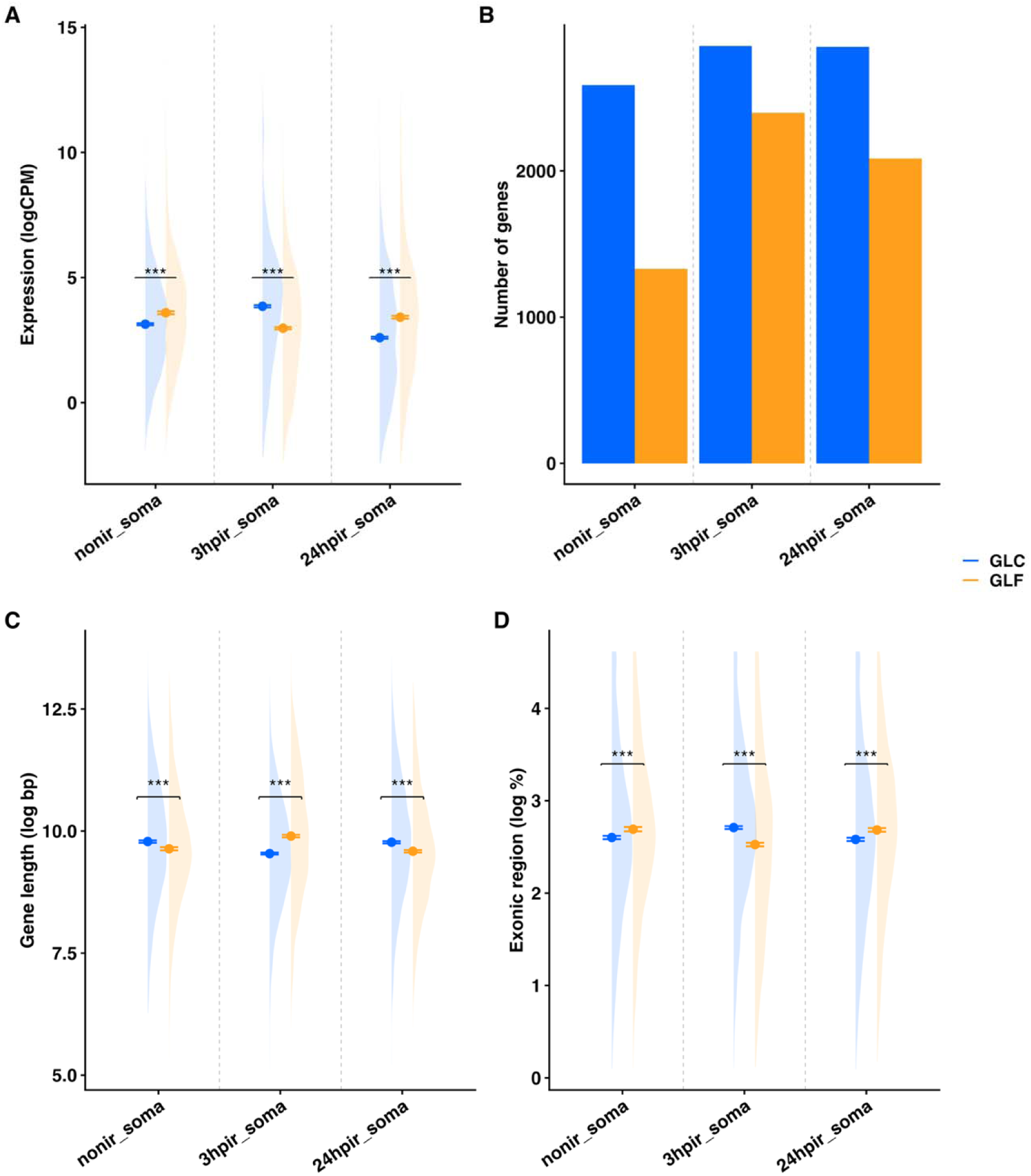
Germline ablation alters the genomic properties of differentially expressed genes in a temporally dynamic manner. Interaction between germline ablation (GLC vs GLF) and irradiation (non-irradiated, 3 hpir, 24 hpir) on somatic gene properties. Contrasts are: nonir_soma (C1), 3hpir_soma (C2.3), and 24hpir_soma (C2.24). **(A)** Mean gene expression (logCPM). **(B)** Number of differentially expressed genes. **(C)** Gene length (bp). **(D)** Percentage exonic sequence. Points denote group means; error bars represent standard errors.

### Baseline differences in DREAM complex architecture

The conserved DREAM repressor complex regulates cell-cycle and DNA repair genes (e.g., *pcna*, *ccna2*, *rad51*, *cdk1*) by silencing their expression during quiescence and after DNA damage^18^. In non-irradiated soma, GLC fish expressed three core components (*lin54*, *rbl1*, *e2f5*), whereas GLF fish expressed *e2f4* only. In addition, GLF fish exhibited upregulated DREAM targets (*pcna*, *ccna2*, *rad51*), suggesting reduced DREAM repression and greater baseline engagement of replication and repair pathways in GLF soma^19^ (Figure 3; Supplementary Figure S7).

**Figure 3.**
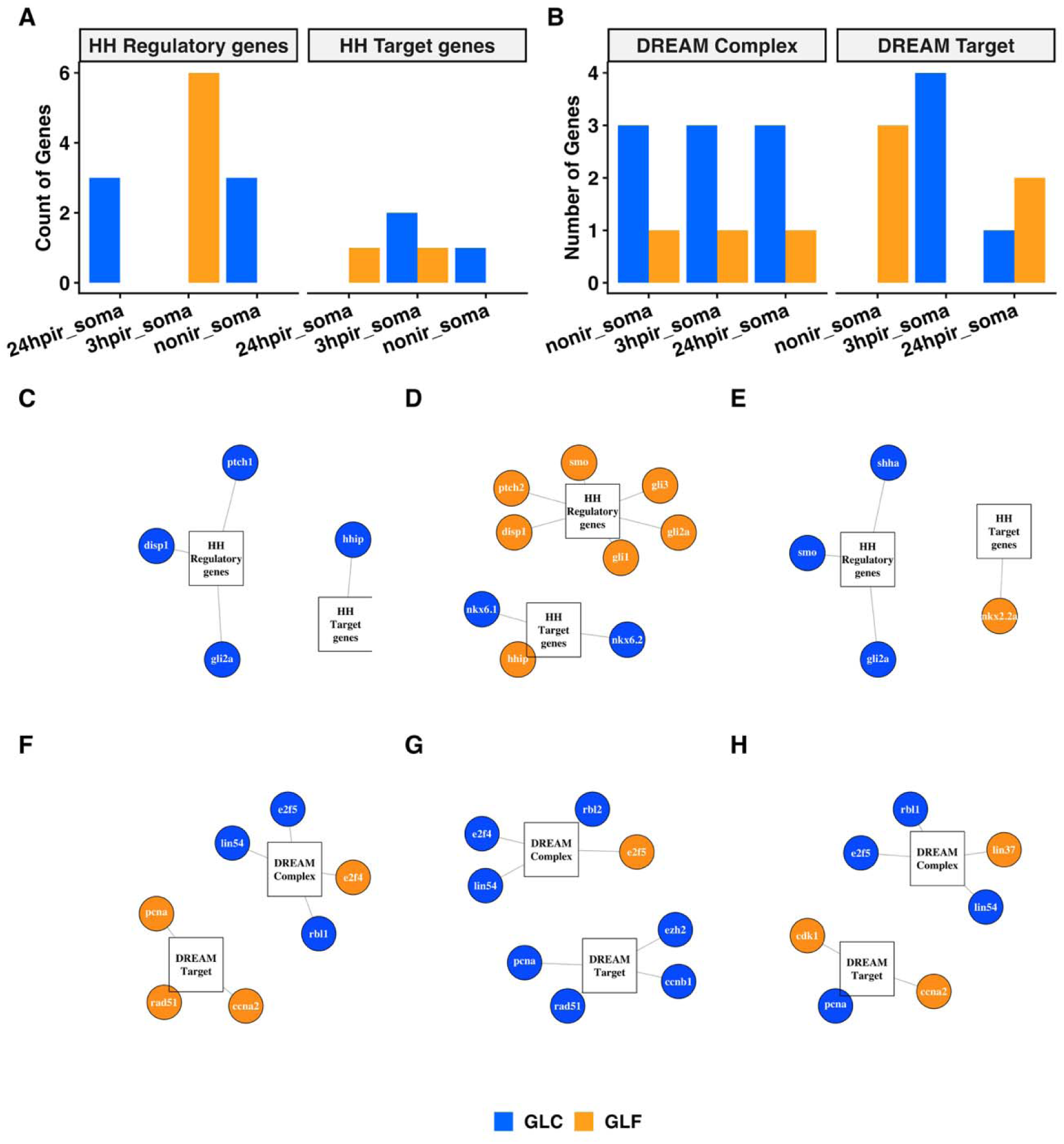
Strain-specific dynamics of DREAM complex and Hedgehog pathway gene expression across irradiation timepoints. Line plots of the number of differentially expressed genes (FDR < 0.05) across timepoints (non-irradiated, 3 hpir, 24 hpir) for **(A)** Hedgehog and **(B)** DREAM complex pathway components and targets, coloured by strain. Network visualisations of DE gene interactions for Hedgehog **(C-E)** and DREAM **(F-H)** pathways at each timepoint: **(C, F)** non-irradiated soma, **(D, G)** 3 hpir, **(E, H)** 24 hpir. Circles represent individual genes; squares represent gene-type categories. Node colour indicates strain. Edges connect genes to their functional category. Networks arranged using the Fruchterman-Reingold layout algorithm.

### Germline ablation accelerates and redirects the somatic response to genotoxic stress

Both strains mounted strong transcriptional responses to 20 Gy γ-irradiation, but with distinct timing and composition (Figure 2). At 3 hpir, 8,252 DEGs were detected (3,779 upregulated in GLF; 4,473 in GLC; Supplementary Table S2), and at 24 hpir, 7,991 DEGs were detected (3,372 upregulated in GLF; 4,619 in GLC). GO term analysis identified substantial shared terms between strains (208 at 3 hpir; 262 at 24 hpir) yet with pronounced divergence in underlying gene composition, especially at 3 hpir (Supplementary Figures S2, S3). Early shared responses concentrated on chromosome segregation, double-strand break (DSB) repair, and RNA catabolism; minimal similarity was observed for protein autophosphorylation, nuclear localisation, and NF-κB signalling. By 24 hpir, similarity increased for ribosomal large subunit biogenesis and RNA metabolism, whereas DNA repair, chromatin remodelling, and nucleocytoplasmic trafficking remained divergent (Figure 2; Supplementary Figure S3).

Among highly divergent functional categories, the “ageing” GO term showed strong temporal and strain dependence. It was supported exclusively by GLC genes at baseline, became shared between strains immediately after irradiation, and by 24 hpir reverted to predominant GLC representation (Figure 4). This pattern suggests that irradiation transiently synchronises ageing-related transcriptional pathways before reinstating strain-specific control.

**Figure 4.**
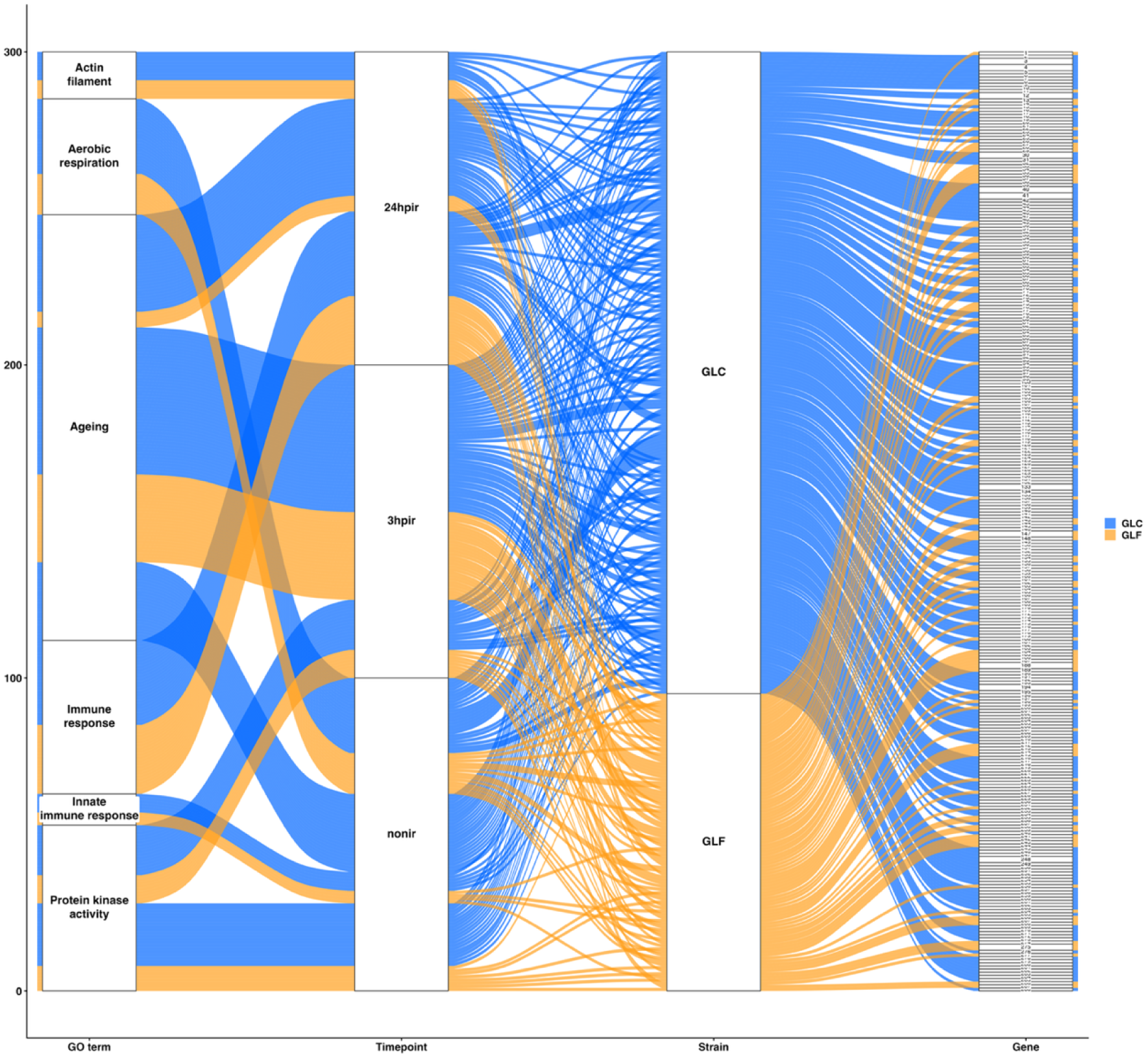
Treatment-dependent conservation of ageing-related GO terms across zebrafish strains. Alluvial plot illustrating relationships between enriched Gene Ontology (GO) terms, experimental condition, strain, and contributing gene sets. Each stratum on the left represents a GO biological process term; flows track associations across irradiation condition/timepoint, strain, and underlying gene sets. Ribbon widths are proportional to the number of contributing genes. Colours denote strain-level clustering. While major ageing-related GO terms are observed in both strains, GLC fish show stronger representation and higher gene-level contribution to ageing- associated processes compared with GLF.

Functionally, the early (3 hpir) response in GLC fish emphasised cellular housekeeping (cytoplasmic translation, ribosome biogenesis), whereas GLF fish exhibited upregulated signal transduction, immune modulation, and structural remodelling (Supplementary Figure S3), indicating a faster adaptive response. By 24 hpir, GLC fish remained skewed towards transcriptional and biosynthetic processes, while GLF fish shifted strongly to DNA replication, mitosis, homologous recombination, and proteasomal degradation (Supplementary Figure S3).

### Temporal reversal in gene expression after irradiation

DEG properties-gene length, exonic density, and expression level-exhibited temporal reversals across irradiation timepoints (Supplementary Tables S6–S7; Figures 1–2). Relative to GLF fish at baseline, GLC fish showed lower expression and exonic density with longer genes; at 3 hpir these differences reversed; and by 24 hpir they reverted towards baseline. Brain and kidney showed the highest divergence in overall gene expression; brain, skin, and spleen showed the highest divergence in gene length; and brain and intestine showed the highest divergence in exonic density.

Genes switching from higher expression in GLF at 3 hpir to higher expression in GLC at 24 hpir were enriched for tissue regeneration, muscle maintenance, and cellular catabolism, supporting an earlier reparative and metabolic reorganisation in GLF^20^. Top DEGs exhibiting this temporal inversion included MHC class II haplotypes (*mhc2dgb*, *mhc2dhb*, *mhc2dga*), which showed similar temporal trends across organs, whereas *mhc2dab* was uniquely elevated in GLC soma (Figures 1–2; Supplementary Table S5).

### Organ-specific pathways reveal rapid response to irradiation in GLF fish

At 3 hpir, GLC organs broadly upregulated biosynthetic metabolism (cytoplasmic translation, ribosome biogenesis, DNA replication), whereas GLF organs upregulated tissue repair, immune signalling, and stress response pathways (Supplementary Figure S3). Notably, GLF brain showed enrichment for coagulation and ERK signalling, intestine and kidney for DNA damage response and NF-κB signalling, and skin for hormone-mediated pathways. At 24 hpir, GLF brain, intestine, and spleen upregulated recombinational repair, chromosome segregation, and mitotic division, with caudal fin showing metabolic adaptation (Supplementary Figure S3), whereas GLC organs upregulated mitochondrial energy metabolism, ribosome biogenesis, and RNA splicing (Figure 1).

### Transcriptional response to irradiation in testes

In GLC testes, irradiation induced large and temporally phased transcriptional changes (3 hpir: 6,710 upregulated/9,118 downregulated; 24 hpir: 6,989/7,736; Supplementary Table S2) with widespread reversals between timepoints (Supplementary Figure S4). Testes mounted a stronger and earlier response than soma, upregulating DNA repair, chromatin remodelling, and transcriptional control at 3 hpir while downregulating many somatic biosynthetic pathways (ribosome biogenesis, RNA splicing, mitochondrial translation; Supplementary Table S8). By 24 hpir, testes upregulated ribosome biogenesis, telomere maintenance, mitochondrial gene expression, and DSB repair, and downregulated immune signalling, returning towards homeostasis faster than any somatic organ.

### Temporal control by DREAM and Hedgehog signalling

Irradiation inverted the baseline DREAM target pattern (Figure 3). At baseline, DREAM targets were GLF-biased (*pcna, ccna2, rad51*); at 3 hpir this was reversed, with GLC upregulating four targets (*ezh2, pcna, ccnb1, rad51*) and GLF none. By 24 hpir the pattern had partially reverted, with GLF expressing ccna2 and cdk1 and GLC expressing pcna, consistent with DREAM’s role in modulating DNA repair gene expression after genotoxic stress^18,19^ (Figure 3). Core DREAM components, by contrast, remained predominantly GLC-biased at every timepoint. Hedgehog (Hh) signalling balances stem-cell maintenance, proliferation, and differentiation in adult tissues and sustains regenerative responses^21,22^. In GLF, Hh regulators (*disp1*, *ptch2*, *gli1*, *gli2a*, *gli3*, *smo*) were upregulated exclusively at 3 hpir; no Hh regulators were differentially expressed in GLF at baseline or by 24 hpir, where the differentially expressed regulators (*shha, smo, gli2a*) were instead GLC-biased (Figure 3), indicating a sharply time-restricted activation of Hh signalling in GLF soma.

### Germline ablation shapes somatic transposable element activity

Transposable elements exhibited strong strain- and time-specific behaviour that closely paralleled gene expression dynamics (Figure 5; Supplementary Figure S9). At baseline, 522 TEs were differentially expressed (239 upregulated in GLF; 283 in GLC; Supplementary Table S9). After irradiation, 229 and 256 TEs were differentially expressed at 3 hpir and 24 hpir, respectively. A directional switch mirrored the gene-expression response: GLF reduced TE expression at 3 hpir and increased it at 24 hpir, whereas GLC showed the opposite pattern (Figure 5), indicating that germline ablation alters not only the timing of gene activation but also the temporal modulation of repetitive-element activity.

**Figure 5.**
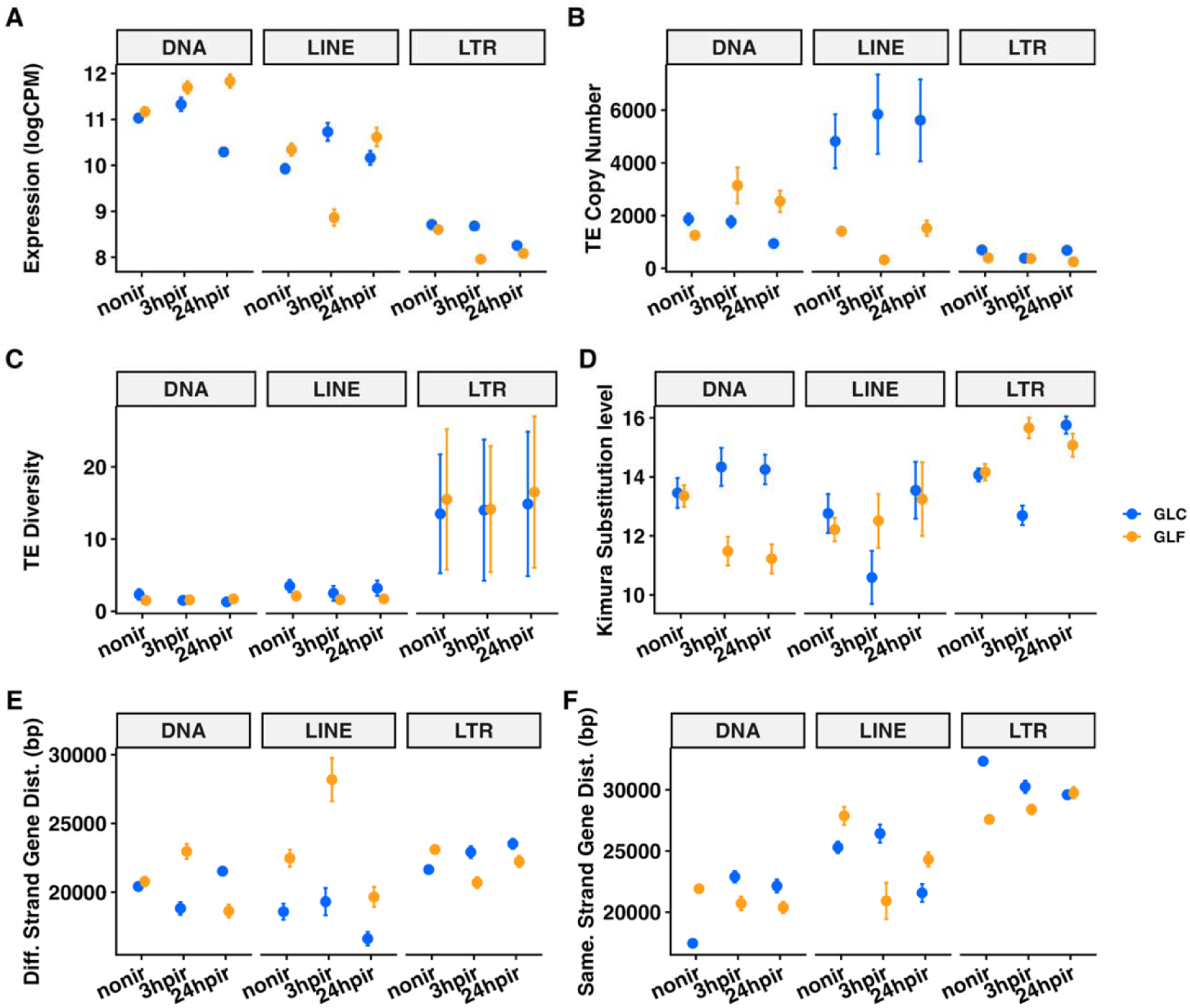
Strain- and timepoint-specific transposable element expression dynamics in somatic tissues following irradiation. **(A)** Mean log counts per million (logCPM) of significantly expressed TE repeats (FDR < 0.05). **(B)** Mean TE copy number per class. **(C)** Number of TE subfamilies differentially expressed. **(D)** Mean synonymous substitution rate (*K*) used as a proxy for TE evolutionary age (lower *K* = more recent insertion). **(E)** Distance of expressed TEs to the nearest gene on the opposite strand (bp). **(F)** Distance to nearest gene on the same strand. Points denote group means; error bars represent standard errors. Strain identity is colour-coded consistently across panels. All panels examine TE metrics across non-irradiated, 3 hpir, and 24 hpir conditions.

At baseline, GLF showed a positive correlation between LINE copy number and diversity. After irradiation, both strains converged on reduced diversity with increased copy number (Supplementary Figure S9). At 3 hpir, GLF LINE expression declined and differentially expressed repeats were located closer to genes on opposite strands, consistent with increased regulatory interference in the absence of the germline. GLC uniquely expressed L2-1_DR at this timepoint. LTR dynamics were largely driven by Gypsy elements and showed an inverse relationship between copy number and diversity across conditions. DNA transposons were particularly elevated in GLF at 24 hpir and were characterised by increased copy number and younger inferred insertion age. At baseline, DE gene-DE TE overlap was greater in GLC (Supplementary Figure S8), suggesting tighter basal coupling between gene expression and TE activity, potentially through shared chromatin accessibility or epigenetic control. After irradiation, overlap increased specifically on chromosome 4 in GLF, dominated by LTR elements^23,24^. No comparable chromosome-wide structure was observed for DE genes or DE TEs in isolation, indicating that the coordinated response emerges from their interaction rather than from independent chromosomal effects.

To investigate whether DE genes are spatially associated with DE TEs, we compared the genomic proximity of DE genes to DE TE loci against all expressed genes as background. For each DE TE family in a given contrast, the single genomic locus closest to any DE gene was retained to avoid inflation from multi-copy TE families. DE genes were consistently enriched within 1–10 kb of DE TE loci compared with all expressed genes (2–3-fold enrichment; Supplementary Figure S10), with enrichment decaying monotonically at larger distances. Among DE gene–TE pairs within 5 kb sharing concordant direction of expression change, LTR retrotransposons were the predominant TE class (Supplementary Figure S10)^23,24^.

## Discussion

At baseline, the soma of GLF fish differentially expressed shorter, exon-rich genes, consistent with reduced transcriptional cost^15–17^ and increased splicing efficiency^16,25^. GLF fish also showed enrichment for cell-cycle progression and chromatin-linked DNA integrity, alongside upregulation of canonical DREAM targets involved in replication and repair^18,26^. Because DREAM represses these targets during quiescence, their elevation indicates a relaxation of cell-cycle brakes in the GLF soma, permitting greater baseline engagement of replication-coupled repair, although we note that the same relaxation removes a tumour-suppressive constraint^27,28^. These results suggest that germline ablation shifts somatic transcription towards a more repair-ready state.

Germline ablation also accelerated the response to genotoxic stress. GLF fish mounted a rapid early response via genes involved in signal transduction, immune modulation, and structural remodelling, transitioning at 24 h to replication-coupled repair and proteostasis^20^. By contrast, GLC fish engaged biosynthetic and transcriptional pathways at both timepoints, delaying their genome maintenance response. The temporal reversals in gene length, exonic density, and expression level across timepoints show that this is a systemic rescheduling of the somatic transcriptional programme rather than an organ-specific effect, and indicate that germline status controls when, not only whether, repair programmes are deployed, which further emphasises the fundamental role the germline plays in somatic repair.

The rescheduling of somatic transcriptional programme is accompanied by changes in two regulatory nodes, DREAM and Hedgehog signalling. The DREAM complex is a conserved master regulator of somatic DNA repair^18^. DREAM target engagement inverted after irradiation in a strain-specific manner, where targets were GLF-biased at baseline, whereas at 3 hpir GLC fish upregulated four targets and GLF fish none, with partial reversion by 24 hpir. By contrast, core DREAM components remained GLC-biased throughout, meaning that the swing in target expression is unlikely to reflect a simple change in repressor abundance. What the data show most clearly is that germline status determines when replication-coupled repair programmes are engaged^18,28^, early and constitutively in germline-free fish, and only under damage in their germline-carrying siblings.

Hedgehog (Hh) signalling, which is vital for tissue regeneration and is dysregulated with age, showed an equally sharp temporal restriction. Hh regulators were differentially expressed in GLF soma at 3 hpir, while at 24 hpir the regulators that differed between strains were instead GLC- biased. Germline-free fish therefore engaged Hh quickly and transiently under genotoxic stress, consistent with the brief, damage-associated activation seen in regenerative contexts^29,30^, rather than with sustained signalling. This is likely important because chronic Hh signalling accelerates ageing, and in *C. elegans*, a piRNA-dependent Hedgehog-related signal travels from germline to soma and actively promotes somatic ageing showed that^31^. Interestingly, no Hh regulators were GLF-biased at baseline, while those differentially expressed in unirradiated fish were GLC-biased. This raises the possibility that the germline sustains basal Hh signalling in the soma, as it does in *C. elegans*^31^, and that germline removal both reduces a chronic pro-ageing signal and restores the capacity for rapid, damage-associated Hh activation.

Germline status also restructured TE activity, and did so in line with gene expression, where the GLF soma suppressed transposable elements early and elevated them at 24 hpir, while the GLC soma showed the opposite pattern. After irradiation, differentially expressed genes and TEs converged specifically on chromosome 4 in GLF fish, with no comparable chromosome-wide structure for either class in isolation, indicating that the coordinated response emerges from their interaction rather than from independent chromosomal effects. This is unlikely to be coincidental because the long arm of zebrafish chromosome 4 is unique in the genome: it is heterochromatic, late-replicating, gene-poor and highly repeat-dense, and is enriched for duplicated zinc-finger gene families^32,33^. This is precisely the chromatin context in which coordinated gene-TE regulation would be expected to concentrate. Moreover, although the zebrafish genome is dominated by type II DNA transposons rather than retroelements^34^, the concordant, proximal pairs we recover are predominantly LTR retrotransposons, which can act as cis-regulatory elements for host stress and immune genes^23,24^. We therefore propose that this reflects germline-dependent regulatory coupling within a permissive chromatin domain, rather than as TE-driven mutagenesis.

Consistent with germline-dependent control of the somatic ageing programme more broadly, the "ageing" GO term was supported exclusively by GLC genes at baseline, was transiently shared after irradiation, and reverted to predominantly GLC representation by 24 hpir. Irradiation therefore appears to synchronise the two strains onto a common ageing-associated programme only briefly, before germline-dependent control is reinstated. The "disposable soma" theory holds that the somatic repair is not maximised because limited resources are allocated to the germline^6,7^, leading to the idea that removing the germline should allow for more resources for the soma. This is not exactly what we observe here. The germline-free soma differentially expressed shorter, cheaper genes, was biased away from biosynthesis under stress in favour of repair and upregulated DNA repair at baseline under reduced DREAM repression. The more parsimonious interpretation is that germline status acts as a regulatory signal for somatic maintenance. This is in line with comparative data in other organisms, for example removing the entire somatic gonad and the germline in *C. elegans* eliminates reproduction without extending lifespan whereas germline removal extends it^34^; similarly, in killifish, germline depletion enhances somatic repair while arrested differentiation does not^4^, and in both *C. remanei*^11^ and *N. furzeri*^4,12^ germline removal extends male but not female lifespan. Recent theoretical work provides a mathematical basis for this argument rooted in Hamiltonian forces of selection^13^. Because suppressed fertility directly strengthens selection against mortality, any condition that reduces reproduction should select for increased somatic maintenance in that condition, with no requirement for resource reallocation or genetic trade-offs. Crucially, this argument is not specific to nutrition, it applies to any stressor that suppresses fertility, and germline damage is one such stressor^13^. Indeed, the model was proposed partly to account for the observation that DNA damage restricted to the germline is sufficient to induce systemic somatic stress resistance in *C. elegans*^5^. Where germline damage transiently suppresses fertility, germline ablation removes it permanently, and the soma we observe, which is repair-ready and faster to respond to stress is what an evolved response to the loss of reproductive prospects should look like. The germline-free soma is therefore not reallocating spared resources but expressing a maintenance programme that is suppressed when the germline is intact. The sex-specificity of germline effects on lifespan^4,11,12^ follows naturally, since the sexes differ in how rapidly selection on survival declines with age^35^, and therefore in how much is gained by expressing that programme.

Our results are transcriptional, and elevated expression of high-fidelity repair genes is a proxy for repair capacity rather than a measurement of repair fidelity; direct assays of repair kinetics and somatic mutation burden are needed to establish that the germline-free soma does in fact protect its genome better. Another limitation is that because *dnd-1* knockout removes primordial germ cells from the outset, germline-derived signalling cannot be fully disentangled from the absence of gametogenesis, as has been done previously in killifish^4^. Finally, we studied males only. Theory predicts that the benefit of germline removal should reflect how rapidly selection on survival declines with age^36^, and therefore that the somatic benefits reported here should be less obvious in females. However, while we did not study females here, the results from earlier studies in other taxa are in line with this prediction because germline removal does not extend female lifespan^4,11,12^.

Our findings identify germline status as a systems-level regulator of somatic genome maintenance and TE dynamics across multiple organs, providing a mechanistic framework for understanding how the germline shapes ageing in vertebrates. More broadly, our results raise the possibility that induction of germline-dependent repair pathways in somatic tissues could be harnessed to improve genome integrity and promote healthy ageing. We predict that manipulating germline-derived signals, rather than somatic resource allocation, will prove the more effective route to enhancing somatic genome stability.

## Materials and Methods

### Fish lines and maintenance

Wild-type AB zebrafish (Danio rerio) were maintained at 28 °C in 3 L tanks under a 12:12 h light– dark cycle at the SciLifeLab, Evolutionary Biology Centre, Uppsala University, in accordance with approved ethical protocols. Water quality (pH 7.0–7.5, conductivity 500–550 μS) was continuously monitored. Juvenile fish were fed five times daily; adults were fed three times daily with live Artemia and dried flake food (Zeigler Adult Zebrafish Diet; Aquatic Habitats). Strict outbreeding was maintained across generations. These fish were then used for CRISPR/Cas9- mediated knockout of *dnd-1* was used to generate germline-free (GLF) males (see supplementary material for details). Heterozygous carriers produced germline-carrying (GLC) siblings. Loss of *dnd-1* results in absence of primordial germ cells, yielding sterile males with normal somatic gonadal development. We used 12 CLC and 12 GLF adult males (>6 months old) for this study.

### Gamma Irradiation

We exposed 12 GLC and 12 GLFmales to 20 Gy dose from a Caesium-137 gamma source (Gammacell 3000; Stockholm University). For exposure, fish were transferred to sterile six-well plates (16.8 mL system water per well, 1–2 fish per well), sealed with laboratory film to prevent desiccation, and irradiated for approximately 25 min. Dosimetry was confirmed with a calibrated ionisation chamber (ISO/IEC 17025 standard). Post-irradiation, fish were returned to standard housing and euthanised and dissected at 3 h and 24 h. All experiments were performed under license no. C28/16 from the Swedish Board of Agriculture.

### Sample collection

We collected five somatic tissues (brain, intestine, kidney, spleen, caudal fin) and testes (GLC only) by dissection from six non-irradiated GLC and GLF males and 6 irradiated GLC and GLF males at 3 hours post irradiation (hpir) and 24 hpir. This resulted in a sample size of three per group. Fish were dissected on ice and organs flash-frozen in liquid nitrogen (Supplementary Figure S1).

### RNA extraction and sequencing

Tissues were manually homogenised in Trizol Reagent (Invitrogen) and RNA extracted following the manufacturer’s protocol. Phase separation was performed with chloroform (centrifugation at 12,000 × g, 20 min, 4 °C). Residual DNA was removed using the TURBO DNA-free Kit (Ambion). RNA concentration and purity were assessed by NanoDrop spectrophotometry; integrity was verified using an Agilent Bioanalyzer. Libraries were prepared using the NEBNext Ultra II Directional RNA Library Prep Kit (poly-A selection, 200–300 bp inserts) and sequenced by Novogene (UK) in two balanced batches (intestine, kidney, and testes: batch 1; brain, spleen, and caudal fin: batch 2) on an Illumina NovaSeq 6000 S1 v1.5 platform (150 bp paired-end, 60-plex per lane).

### RNA-seq processing pipeline

Raw reads were processed using the nf-core RNA-seq pipeline^37^ (v3.9) under nextflow^38^ (v22.04.5). Quality trimming and adapter removal was performed with Trim Galore^39^ (v0.6.10) (reads ≥ 10,000 per sample retained). Reads were aligned to the *Danio rerio* reference Genome (GRCz11) using STAR^40^ (v2.7.11, two-pass mode) and quantified with SALMON^41^ (v1.10.1) in selective-alignment mode. Gene-level count matrices were generated using nf-core standard parameters. Quality metrics were assessed using FastQC ^34^ (v0.12.1) and MultiQC^42^ (v1.14)

### Differential gene expression analysis

Gene-level counts were analysed in edgeR^43^ (v3.38; Bioconductor) in RStudio^44^ v.4.5.3. Genes were retained if CPM > 1 in ≥3 samples. Normalisation used TMM (trimmed mean of M values). Differential expression was tested using generalised linear models (GLMs) with the following contrasts: (i) GLC vs GLF non-irradiated (Contrast 1); (ii) irradiated vs non-irradiated within each strain at 3 hpir (Contrast 2.3) and 24 hpir (Contrast 2.24); and (iii) organ-specific strain × irradiation interactions (Contrast 3– Contrast 5). Germline responses were modelled separately (Contrast 6.3, Contrast 6.24). Details on exact comparisons for each contrast can be found in Supplementary Table S3. Statistical significance was assessed using Benjamini– Hochberg FDR correction at FDR < 0.05. Volcano plots were generated with EnhancedVolcano^45^ (v1.18).

### Gene Ontology and chromatin analysis

GO enrichment was performed using PathfindR^46^ (v2.1.0) with zebrafish annotations from MSigDB. Human orthologues were identified via BiomaRt. Gene coordinates were mapped to ATAC-seq chromatin accessibility tracks from the zebrafish COPES dataset^47^ identifying constitutively open chromatin regions across zebrafish development. Exonic and intronic lengths were calculated from Ensembl annotations (v107; Danio rerio GRCz11). Exonic regions were merged by chromosomal start position to avoid artificial elongation from alternative splicing; total exonic length per gene was computed as the sum of non-overlapping exonic regions. Exonic proportion was expressed as (exonic length / gene length) × 100.

### Transposable element analysis

TE expression was quantified using TEtranscripts^48^ (v2.2.3) with the *Danio rerio* genome (GRCz11) and Ensembl TE annotations. TE differential expression was analysed analogously to gene-level data using edgeR. Because libraries were poly(A)-selected, detected TEs represent polyadenylated fractions of transposon-derived transcripts. TE evolutionary age and divergence were inferred using RepeatMasker^49^ and calcDivergenceFromAlign.pl, which calculate Kimura substitution levels (K) between TE copies and their consensus sequences. Lower K values indicate more recent transposon activity. TE copy-number density was calculated as the proportion of TE-occupied base pairs per gene, normalised by total gene length.

### Data Availability

All RNA-seq data are available on ENA under accession PRJEB102813 (under embargo until acceptance). Scripts and supplementary data are available at: https://github.com/asdfjohnny1/germline-ablation-transcriptomics.git. Detailed bioinformatic pipeline settings are provided in Supplementary Methods.

## Acknowledgements

This research was funded by a Consolidator Grant from the European Research Council (SELECTHAPLOID-101001341) to SI.

## Author Contributions

Conceptualisation: S.I., G.A.; Methodology: J.C.D.C., G.A., E.T., M.S.; Software: J.C.D.C.; Formal analysis: J.C.D.C.; Investigation: G.A., E.T., M.S.; Resources: S.I., R.R.; Data curation: J.C.D.C.; Writing – original draft: J.C.D.C., S.I., A.A.M.; Writing, review & editing: All authors; Visualisation: J.C.D.C.; Supervision: S.I., W.H., R.R., A.A.M.; Project administration: S.I.; Funding acquisition: S.I.; Bioinformatic support: A.M.G.

