## Supplementary information for "Germline removal reprograms somatic genome maintenance towards faster, energy-efficient, high-fidelity DNA defence and repair"

**Supporting Information**

**Germline removal reprograms somatic genome maintenance towards faster, energy-efficient, high-fidelity DNA repair and enhanced transposable element control**

Jean-Charles de Coriolis, Ghazal Alavioon, Wilfried Haerty, Emmanouil Tsakoumis, Monika Schmitz, Alice M. Godden, Raheleh Rahbari, Alexei A. Maklakov, Simone Immler

**This file includes:**

• Supporting text (Supplementary Methods)

• Supplementary Figures S1–S10

• Supplementary Tables S1–S9

### Supplementary Methods

#### *CRISPR/Cas9-Mediated Germline Ablation*

##### sgRNA design and template preparation

Target sequences within *dnd-1* were identified using CRISPR-Scan (https://www.crisprscan.org) and verified for minimal off-target potential. Each 20-nt target sequence was prefixed with the T7 initiation site (“GG”) and appended to a 20-nt overlap complementary to the generic sgRNA scaffold. The targeting oligonucleotide (oligo A) was annealed to a universal sgRNA backbone oligonucleotide (oligo B; 5′-AAAAGCACCGACTCGGTGCCACTTTTTCAAGTTGATAACGGACTAGCCTTATTTTAACTTGCTATTTCTAGCTCTAAAAC-3′) following Varshney et al. (2015). Annealing and fill-in reactions used Phusion High-Fidelity DNA Polymerase (NEB M0530S): 98 °C for 2 min, 50 °C for 10 min, 72 °C for 10 min. The expected 120 bp sgRNA template was confirmed by electrophoresis on a 2.5% agarose gel.

##### In vitro transcription of sgRNA

Approximately 2–3 μL of assembled sgRNA template was used as input for RNA synthesis using the T7 Quick High Yield RNA Synthesis Kit (NEB E2050S). Following transcription, reactions were treated with DNase I at 37 °C for 15 min, and RNA was purified by isopropanol/sodium acetate precipitation or using the mirVana miRNA Isolation Kit (Life Technologies AM1560). RNA was quantified by NanoDrop (expected yield 1.5–3 μg/μL) and quality-checked on a 2–2.5% agarose gel. Aliquots were stored at −80 °C.

##### Preparation of Cas9 mRNA

Cas9 mRNA was synthesised from pT3TS-nCas9n (Addgene #46757), linearised with XbaI (NEB) and purified using a QIAprep Spin Column (Qiagen 27104). In vitro transcription used the mMESSAGE mMACHINE T3 Kit (Life Technologies AM1348). RNA was purified by LiCl precipitation or RNeasy Mini Kit (Qiagen 74104), integrity verified on a 1% agarose gel (expected single band ~4.5 kb), and stored at −80 °C (>1 μg/μL).

##### Microinjection

Injection mixtures were prepared on ice: 1 μL Cas9 mRNA (750 ng/μL), 1 μL sgRNA per target (250 ng/μL), 1 μL phenol red (Sigma P0290), and RNase-free water to 10 μL total. Needles were calibrated to deliver 2 nL per embryo (~150 pg Cas9 mRNA; ~25 pg sgRNA per target). Injections were performed at the one-cell stage into the yolk (higher embryo survival) or cell (greater mutagenesis efficiency). Embryos were maintained at 28 °C in E3 medium and inspected daily.

##### Verification of mutagenesis and germline loss

At 2–3 days post-fertilisation, embryo subsets were screened for editing efficiency using the T7 Endonuclease I assay (NEB M0302L) or fragment length analysis. Successfully edited fish were raised to adulthood and confirmed to exhibit complete germ cell ablation by morphological and genotypic analysis.

#### *Spatial Association Between Genes and Transposable Elements*

Genomic coordinates for DE TE families were retrieved from the UCSC RepeatMasker annotation for Danio rerio (GRCz11) via AnnotationHub (Bioconductor). For each contrast × strain context, the single RepeatMasker locus per DE TE family name closest to any DE gene was identified using distanceToNearest (GenomicRanges v1.50; ignore.strand = TRUE), retaining one locus per TE family to avoid inflation from high-copy-number families. Fold enrichment was calculated as the proportion of DE genes within each distance threshold (1, 5, 10, 50, 100, 500 kb) divided by the corresponding proportion for all expressed genes in that context. Concordant gene–TE pairs were defined as pairs within 5 kb where the gene and TE showed the same direction of fold change in the same contrast × strain context. Statistical significance of DE gene proximity enrichment over background was assessed using one-sided Wilcoxon rank-sum tests with Benjamini–Hochberg correction.

#### *Mixed-Effects Modelling of Gene Structural Properties*

Linear mixed-effects models (LMMs) were fitted in R using lme4 (lmer) to analyse gene expression (logCPM) and gene length, with fixed effects of strain, contrast, gene length (as covariate), and chromatin openness (COPES), and random effects of chromosome and strand. Generalised linear mixed-effects models (GLMMs; glmer with binomial family) were applied to the proportion of exonic sequence. Significance of fixed effects was assessed by likelihood ratio tests (χ²). Pairwise comparisons of estimated marginal means were derived using the emmeans package with Tukey HSD correction.

### Supplementary Tables

#### Table S1. Generalised linear model specifications. Two GLMs were fitted in edgeR to biologically defined sample subsets. Model dispersion (global overdispersion across genes), mean tagwise dispersion (gene-level variability), and residual degrees of freedom are reported.

| **Model** | **edgeR formula** | **Comparison** | **N samples** | **Dispersion** | **Mean tagwise dispersion** | **Residual df** |
| --- | --- | --- | --- | --- | --- | --- |
| Full soma | ~ Strain × Irradiation × Tissue × Timepoint + Individual | Strain × irradiation effects across somatic organs | 111 / 123 | 2.4 | 3.7 | 55 |
| Germline | ~ Irradiation × Timepoint + Individual | Effect of irradiation on germline gene expression | 12 / 123 | 0.6 | 1.2 | 8 |

#### Table S2. Number of differentially expressed genes per contrast (FDR < 0.05). Upregulated genes have logFC > 0; downregulated genes have logFC < 0. Contrast labels: C1, non-irradiated soma; C2.3/C2.24, irradiated soma at 3/24 hpir; C3, organ-specific baseline; C4/C5, organ-specific 3/24 hpir; C6.3/C6.24, testes at 3/24 hpir.

| **Contrast** | **Upregulated** | **Downregulated** | **Non-DE** | **Total expressed** |
| --- | --- | --- | --- | --- |
| C1 | 2,357 | 4,498 | 16,659 | 23,514 |
| C2.3 | 3,779 | 4,473 | 15,262 | 23,514 |
| C2.24 | 3,372 | 4,619 | 15,523 | 23,514 |
| C3.brain | 1,592 | 3,891 | 18,031 | 23,514 |
| C3.intestine | 3,211 | 6,476 | 13,827 | 23,514 |
| C3.kidney | 3,405 | 6,823 | 13,286 | 23,514 |
| C3.caudal fin | 5,990 | 3,766 | 13,758 | 23,514 |
| C3.spleen | 3,850 | 5,692 | 13,972 | 23,514 |
| C4.brain.3 | 4,498 | 4,110 | 14,906 | 23,514 |
| C4.intestine.3 | 5,188 | 4,688 | 13,638 | 23,514 |
| C4.kidney.3 | 4,997 | 4,759 | 13,758 | 23,514 |
| C4.caudal fin.3 | 4,806 | 4,654 | 14,054 | 23,514 |
| C4.spleen.3 | 4,819 | 6,876 | 11,819 | 23,514 |
| C5.brain.24 | 1,884 | 3,050 | 18,580 | 23,514 |
| C5.intestine.24 | 4,465 | 7,002 | 12,047 | 23,514 |
| C5.kidney.24 | 6,117 | 5,270 | 12,127 | 23,514 |
| C5.caudal fin.24 | 5,629 | 5,516 | 12,369 | 23,514 |
| C5.spleen.24 | 4,122 | 4,024 | 15,368 | 23,514 |
| C6.3 (testes) | 6,710 | 9,118 | 7,064 | 22,892 |
| C6.24 (testes) | 6,989 | 7,736 | 8,167 | 22,892 |

#### Table S3. Linear contrasts derived from the full model.

| **Model** | **Contrast ID** | **Comparison** | **Interpretation** |
| --- | --- | --- | --- |
| Soma | C1 | Non-irrad. GLC vs GLF | Effect of germline ablation on non-irradiated soma |
| Soma | C2.3 | 3hpir GLC vs GLF | Germline–irradiation interaction at 3 hpir |
| Soma | C2.24 | 24hpir GLC vs GLF | Germline–irradiation interaction at 24 hpir |
| Soma | C3.brain | Non-irrad. GLC brain vs GLF brain | Germline ablation effect in brain |
| Soma | C3.intestine | Non-irrad. GLC intestine vs GLF intestine | Germline ablation effect in intestine |
| Soma | C3.kidney | Non-irrad. GLC kidney vs GLF kidney | Germline ablation effect in kidney |
| Soma | C3.caudal fin | Non-irrad. GLC caudal fin vs GLF caudal fin | Germline ablation effect in caudal fin |
| Soma | C3.spleen | Non-irrad. GLC spleen vs GLF spleen | Germline ablation effect in spleen |
| Soma | C4.brain.3 | 3hpir GLC brain vs GLF brain | Germline–irradiation interaction in brain at 3 hpir |
| Soma | C4.intestine.3 | 3hpir GLC intestine vs GLF intestine | Germline–irradiation interaction in intestine at 3 hpir |
| Soma | C4.kidney.3 | 3hpir GLC kidney vs GLF kidney | Germline–irradiation interaction in kidney at 3 hpir |
| Soma | C4.caudal fin.3 | 3hpir GLC caudal fin vs GLF caudal fin | Germline–irradiation interaction in caudal fin at 3 hpir |
| Soma | C4.spleen.3 | 3hpir GLC spleen vs GLF spleen | Germline–irradiation interaction in spleen at 3 hpir |
| Soma | C5.brain.24 | 24hpir GLC brain vs GLF brain | Germline–irradiation interaction in brain at 24 hpir |
| Soma | C5.intestine.24 | 24hpir GLC intestine vs GLF intestine | Germline–irradiation interaction in intestine at 24 hpir |
| Soma | C5.kidney.24 | 24hpir GLC kidney vs GLF kidney | Germline–irradiation interaction in kidney at 24 hpir |
| Soma | C5.caudal fin.24 | 24hpir GLC caudal fin vs GLF caudal fin | Germline–irradiation interaction in caudal fin at 24 hpir |
| Soma | C5.spleen.24 | 24hpir GLC spleen vs GLF spleen | Germline–irradiation interaction in spleen at 24 hpir |
| Germline | C6.3 | Non-irrad. GLC testes vs 3hpir GLC testes | Irradiation effect in germline at 3 hpir |
| Germline | C6.24 | Non-irrad. GLC testes vs 24hpir GLC testes | Irradiation effect in germline at 24 hpir |

#### Table S4. Top differentially expressed genes in the soma across irradiation timepoints.Top ten DE genes (ranked by FDR) for each strain (GLC, GLF) at non-irradiated, 3 hpir, and 24 hpir timepoints in the soma. Ensembl gene identifiers, gene symbols, model labels, strain, and primary GO terms are provided. See main text for gene descriptions. Full data available at the project GitHub repository.

| Timepoint | **Ensemble gene ID** | **Symbol** | Model | **Label** | **Strain** | **GO ID** |
| --- | --- | --- | --- | --- | --- | --- |
| Non-irradiated controls (nonir) | ENSDARG00000079105 | mhc2dab | full_model | nonir_soma | GLC | GO:0042613 |
|  | ENSDARG00000097137 | NA | full_model | nonir_soma | GLC |  |
|  | ENSDARG00000071024 | zgc:171679 | full_model | nonir_soma | GLC | GO:0005515 |
|  | ENSDARG00000030448 | sppl2 | full_model | nonir_soma | GLC | GO:0042500 |
|  | ENSDARG00000044074 | loxl2b | full_model | nonir_soma | GLC | GO:0016641 |
|  | ENSDARG00000086418 | si:ch211-236p5.3 | full_model | nonir_soma | GLF | GO:0005515 |
|  | ENSDARG00000087299 | armc9 | full_model | nonir_soma | GLF | GO:0036064 |
|  | ENSDARG00000112702 | NA | full_model | nonir_soma | GLF |  |
|  | ENSDARG00000068621 | nlrc11 | full_model | nonir_soma | GLF | GO:0005515 |
|  | ENSDARG00000104899 | si:dkey-242k1.4 | full_model | nonir_soma | GLF |  |
| 3 hours post-irradiation (3hpir) | ENSDARG00000071024 | zgc:171679 | full_model | 3hpir_soma | GLC | GO:0005515 |
|  | ENSDARG00000003132 | apip | full_model | 3hpir_soma | GLC |  |
|  | ENSDARG00000097137 | NA | full_model | 3hpir_soma | GLC |  |
|  | ENSDARG00000098030 | NA | full_model | 3hpir_soma | GLC |  |
|  | ENSDARG00000103841 | NA | full_model | 3hpir_soma | GLC | GO:0003824 |
|  | ENSDARG00000068621 | nlrc11 | full_model | 3hpir_soma | GLF | GO:0005515 |
|  | ENSDARG00000104635 | mhc2dgb | full_model | 3hpir_soma | GLF |  |
|  | ENSDARG00000091847 | LOC110437747 | full_model | 3hpir_soma | GLF | GO:0005515 |
|  | ENSDARG00000116524 | mhc2dhb | full_model | 3hpir_soma | GLF | GO:0042613 |
|  | ENSDARG00000079645 | NA | full_model | 3hpir_soma | GLF |  |
| 24 hours post -irradiation (24hpir) | ENSDARG00000079105 | mhc2dab | full_model | 24hpir_soma | GLC | GO:0042613 |
|  | ENSDARG00000104635 | mhc2dgb | full_model | 24hpir_soma | GLC |  |
|  | ENSDARG00000071024 | zgc:171679 | full_model | 24hpir_soma | GLC | GO:0005515 |
|  | ENSDARG00000044074 | loxl2b | full_model | 24hpir_soma | GLC | GO:0016641 |
|  | ENSDARG00000074653 | si:ch211-233m11.1 | full_model | 24hpir_soma | GLC | GO:0005515 |
|  | ENSDARG00000068621 | nlrc11 | full_model | 24hpir_soma | GLF | GO:0005515 |
|  | ENSDARG00000091847 | LOC110437747 | full_model | 24hpir_soma | GLF | GO:0005515 |
|  | ENSDARG00000094210 | fthl31 | full_model | 24hpir_soma | GLF | GO:0006879 |
|  | ENSDARG00000087299 | armc9 | full_model | 24hpir_soma | GLF | GO:0036064 |
|  | ENSDARG00000104613 | NA | full_model | 24hpir_soma | GLF |  |

#### Table S5. Top differentially expressed genes per somatic tissue and irradiation timepoint.

Top ten DE genes per tissue (brain, intestine, kidney, caudal fin, spleen) and timepoint for each strain, with Ensembl identifiers and primary GO annotations. Full data available at the project GitHub repository.

| Timepoint | **Ensemble gene ID** | **Symbol** | Model | **Label** | **Strain** | **GO ID** |
| --- | --- | --- | --- | --- | --- | --- |
| Non-irradiated controls (nonir) | ENSDARG00000079105 | mhc2dab | full_model | nonir_brain | GLC | GO:0042613 |
|  | ENSDARG00000044074 | loxl2b | full_model | nonir_brain | GLC | GO:0016641 |
|  | ENSDARG00000097137 | NA | full_model | nonir_brain | GLC |  |
|  | ENSDARG00000113971 | LOC100149352 | full_model | nonir_brain | GLC | GO:0005634 |
|  | ENSDARG00000004141 | dhrs11b.2 | full_model | nonir_brain | GLC | GO:0005575 |
|  | ENSDARG00000087299 | armc9 | full_model | nonir_brain | GLF | GO:0036064 |
|  | ENSDARG00000112702 | NA | full_model | nonir_brain | GLF |  |
|  | ENSDARG00000086418 | si:ch211-236p5.3 | full_model | nonir_brain | GLF | GO:0005515 |
|  | ENSDARG00000027992 | hao2 | full_model | nonir_brain | GLF | GO:0016491 |
|  | ENSDARG00000095949 | si:dkey-22i16.9 | full_model | nonir_brain | GLF |  |
|  | ENSDARG00000079105 | mhc2dab | full_model | nonir_intestine | GLC | GO:0042613 |
|  | ENSDARG00000104387 | slc4a5b | full_model | nonir_intestine | GLC | GO:0008509 |
|  | ENSDARG00000097137 | NA | full_model | nonir_intestine | GLC |  |
|  | ENSDARG00000087012 | NA | full_model | nonir_intestine | GLC | GO:0005525 |
|  | ENSDARG00000036045 | penkb | full_model | nonir_intestine | GLC | GO:0007218 |
|  | ENSDARG00000112702 | NA | full_model | nonir_intestine | GLF |  |
|  | ENSDARG00000076043 | si:dkeyp-73d8.9 | full_model | nonir_intestine | GLF |  |
|  | ENSDARG00000105731 | LOC103908715 | full_model | nonir_intestine | GLF | GO:0005515 |
|  | ENSDARG00000086418 | si:ch211-236p5.3 | full_model | nonir_intestine | GLF | GO:0005515 |
|  | ENSDARG00000078844 | LOC100007883 | full_model | nonir_intestine | GLF |  |
|  | ENSDARG00000079105 | mhc2dab | full_model | nonir_kidney | GLC | GO:0042613 |
|  | ENSDARG00000038587 | LOC100004199 | full_model | nonir_kidney | GLC |  |
|  | ENSDARG00000078828 | npb | full_model | nonir_kidney | GLC | GO:0001664 |
|  | ENSDARG00000055240 | xdh | full_model | nonir_kidney | GLC | GO:0043546 |
|  | ENSDARG00000097137 | NA | full_model | nonir_kidney | GLC |  |
|  | ENSDARG00000112702 | NA | full_model | nonir_kidney | GLF |  |
|  | ENSDARG00000100851 | gphnb | full_model | nonir_kidney | GLF | GO:0032324 |
|  | ENSDARG00000086418 | si:ch211-236p5.3 | full_model | nonir_kidney | GLF | GO:0005515 |
|  | ENSDARG00000086569 | zgc:172051 | full_model | nonir_kidney | GLF |  |
|  | ENSDARG00000035694 | stm | full_model | nonir_kidney | GLF |  |
|  | ENSDARG00000079105 | mhc2dab | full_model | nonir_caudal fin | GLC | GO:0042613 |
|  | ENSDARG00000097137 | NA | full_model | nonir_caudal fin | GLC |  |
|  | ENSDARG00000088625 | si:ch211-142d6.2 | full_model | nonir_caudal fin | GLC | GO:0005515 |
|  | ENSDARG00000113971 | LOC100149352 | full_model | nonir_caudal fin | GLC | GO:0005634 |
|  | ENSDARG00000030448 | sppl2 | full_model | nonir_caudal fin | GLC | GO:0042500 |
|  | ENSDARG00000086418 | si:ch211-236p5.3 | full_model | nonir_caudal fin | GLF | GO:0005515 |
|  | ENSDARG00000087299 | armc9 | full_model | nonir_caudal fin | GLF | GO:0036064 |
|  | ENSDARG00000116832 | NA | full_model | nonir_caudal fin | GLF | GO:0015074 |
|  | ENSDARG00000043002 | vmo1a | full_model | nonir_caudal fin | GLF |  |
|  | ENSDARG00000037789 | pvalb1 | full_model | nonir_caudal fin | GLF | GO:0005509 |
|  | ENSDARG00000079105 | mhc2dab | full_model | nonir_spleen | GLC | GO:0042613 |
|  | ENSDARG00000038587 | LOC100004199 | full_model | nonir_spleen | GLC |  |
|  | ENSDARG00000030448 | sppl2 | full_model | nonir_spleen | GLC | GO:0042500 |
|  | ENSDARG00000031952 | mb | full_model | nonir_spleen | GLC | GO:0015671 |
|  | ENSDARG00000097137 | NA | full_model | nonir_spleen | GLC |  |
|  | ENSDARG00000020084 | tg | full_model | nonir_spleen | GLF |  |
|  | ENSDARG00000086418 | si:ch211-236p5.3 | full_model | nonir_spleen | GLF | GO:0005515 |
|  | ENSDARG00000028148 | pax2a | full_model | nonir_spleen | GLF | GO:0006355 |
|  | ENSDARG00000005392 | slc5a5 | full_model | nonir_spleen | GLF | GO:0055085 |
|  | ENSDARG00000069296 | moxd1l | full_model | nonir_spleen | GLF | GO:0004500 |
| 3 hours post-irradiation (3hpir) | ENSDARG00000114577 | LOC108183819 | full_model | 3hpir_brain | GLC | GO:0005515 |
|  | ENSDARG00000044074 | loxl2b | full_model | 3hpir_brain | GLC | GO:0016641 |
|  | ENSDARG00000003132 | apip | full_model | 3hpir_brain | GLC |  |
|  | ENSDARG00000098030 | NA | full_model | 3hpir_brain | GLC |  |
|  | ENSDARG00000097137 | NA | full_model | 3hpir_brain | GLC |  |
|  | ENSDARG00000068621 | nlrc11 | full_model | 3hpir_brain | GLF | GO:0005515 |
|  | ENSDARG00000104635 | mhc2dgb | full_model | 3hpir_brain | GLF |  |
|  | ENSDARG00000097610 | NA | full_model | 3hpir_brain | GLF |  |
|  | ENSDARG00000116524 | mhc2dhb | full_model | 3hpir_brain | GLF | GO:0042613 |
|  | ENSDARG00000091847 | LOC110437747 | full_model | 3hpir_brain | GLF | GO:0005515 |
|  | ENSDARG00000088524 | acot17 | full_model | 3hpir_intestine | GLC | GO:0016790 |
|  | ENSDARG00000104387 | slc4a5b | full_model | 3hpir_intestine | GLC | GO:0008509 |
|  | ENSDARG00000077068 | si:ch211-11p18.6 | full_model | 3hpir_intestine | GLC | GO:0005515 |
|  | ENSDARG00000016623 | si:ch211-195b13.1 | full_model | 3hpir_intestine | GLC |  |
|  | ENSDARG00000087012 | NA | full_model | 3hpir_intestine | GLC | GO:0005525 |
|  | ENSDARG00000104635 | mhc2dgb | full_model | 3hpir_intestine | GLF |  |
|  | ENSDARG00000071216 | si:ch211-133n4.9 | full_model | 3hpir_intestine | GLF | GO:0003676 |
|  | ENSDARG00000092833 | si:dkeyp-1h4.8 | full_model | 3hpir_intestine | GLF |  |
|  | ENSDARG00000079105 | mhc2dab | full_model | 3hpir_intestine | GLF | GO:0042613 |
|  | ENSDARG00000094210 | fthl31 | full_model | 3hpir_intestine | GLF | GO:0006879 |
|  | ENSDARG00000071223 | zgc:158445 | full_model | 3hpir_kidney | GLC | GO:0005575 |
|  | ENSDARG00000097137 | NA | full_model | 3hpir_kidney | GLC |  |
|  | ENSDARG00000003132 | apip | full_model | 3hpir_kidney | GLC |  |
|  | ENSDARG00000036846 | anks4b | full_model | 3hpir_kidney | GLC | GO:0005515 |
|  | ENSDARG00000078828 | npb | full_model | 3hpir_kidney | GLC | GO:0001664 |
|  | ENSDARG00000091847 | LOC110437747 | full_model | 3hpir_kidney | GLF | GO:0005515 |
|  | ENSDARG00000104635 | mhc2dgb | full_model | 3hpir_kidney | GLF |  |
|  | ENSDARG00000068621 | nlrc11 | full_model | 3hpir_kidney | GLF | GO:0005515 |
|  | ENSDARG00000078529 | adgrb1b | full_model | 3hpir_kidney | GLF |  |
|  | ENSDARG00000079645 | NA | full_model | 3hpir_kidney | GLF |  |
|  | ENSDARG00000042988 | slc24a2 | full_model | 3hpir_caudal fin | GLC |  |
|  | ENSDARG00000098030 | NA | full_model | 3hpir_caudal fin | GLC |  |
|  | ENSDARG00000097480 | NA | full_model | 3hpir_caudal fin | GLC |  |
|  | ENSDARG00000025783 | si:ch211-125e6.11 | full_model | 3hpir_caudal fin | GLC |  |
|  | ENSDARG00000074754 | ifi27.6 | full_model | 3hpir_caudal fin | GLC | GO:0016020 |
|  | ENSDARG00000068621 | nlrc11 | full_model | 3hpir_caudal fin | GLF | GO:0005515 |
|  | ENSDARG00000104635 | mhc2dgb | full_model | 3hpir_caudal fin | GLF |  |
|  | ENSDARG00000103716 | mhc2dga | full_model | 3hpir_caudal fin | GLF | GO:0042613 |
|  | ENSDARG00000024829 | tnn | full_model | 3hpir_caudal fin | GLF | GO:0005515 |
|  | ENSDARG00000116524 | mhc2dhb | full_model | 3hpir_caudal fin | GLF | GO:0042613 |
|  | ENSDARG00000016494 | ddc | full_model | 3hpir_spleen | GLC |  |
|  | ENSDARG00000071601 | pvalb9 | full_model | 3hpir_spleen | GLC | GO:0005509 |
|  | ENSDARG00000068372 | zgc:158427 | full_model | 3hpir_spleen | GLC | GO:0016020 |
|  | ENSDARG00000057789 | lyz | full_model | 3hpir_spleen | GLC | GO:0003796 |
|  | ENSDARG00000088524 | acot17 | full_model | 3hpir_spleen | GLC | GO:0016790 |
|  | ENSDARG00000020084 | tg | full_model | 3hpir_spleen | GLF |  |
|  | ENSDARG00000104635 | mhc2dgb | full_model | 3hpir_spleen | GLF |  |
|  | ENSDARG00000068621 | nlrc11 | full_model | 3hpir_spleen | GLF | GO:0005515 |
|  | ENSDARG00000103716 | mhc2dga | full_model | 3hpir_spleen | GLF | GO:0042613 |
|  | ENSDARG00000116524 | mhc2dhb | full_model | 3hpir_spleen | GLF | GO:0042613 |
|  | ENSDARG00000078683 | LOC110437942 | germline | 3hpir_testes | GLC | GO:0016567 |
|  | ENSDARG00000115978 | NA | germline | 3hpir_testes | GLC |  |
|  | ENSDARG00000075352 | brinp3b | germline | 3hpir_testes | GLC | GO:0005737 |
|  | ENSDARG00000032631 | ltb4r | germline | 3hpir_testes | GLC |  |
|  | ENSDARG00000069282 | bbc3 | germline | 3hpir_testes | GLC | GO:0090200 |
| 24 hours post-irradiation (3hpir) | ENSDARG00000079105 | mhc2dab | full_model | 24hpir_brain | GLC | GO:0042613 |
|  | ENSDARG00000044074 | loxl2b | full_model | 24hpir_brain | GLC | GO:0016641 |
|  | ENSDARG00000104635 | mhc2dgb | full_model | 24hpir_brain | GLC |  |
|  | ENSDARG00000037954 | tnnt1 | full_model | 24hpir_brain | GLC | GO:0005861 |
|  | ENSDARG00000091280 | si:ch211-66k16.27 | full_model | 24hpir_brain | GLC | GO:0042981 |
|  | ENSDARG00000068621 | nlrc11 | full_model | 24hpir_brain | GLF | GO:0005515 |
|  | ENSDARG00000091847 | LOC110437747 | full_model | 24hpir_brain | GLF | GO:0005515 |
|  | ENSDARG00000076043 | si:dkeyp-73d8.9 | full_model | 24hpir_brain | GLF |  |
|  | ENSDARG00000087299 | armc9 | full_model | 24hpir_brain | GLF | GO:0036064 |
|  | ENSDARG00000086418 | si:ch211-236p5.3 | full_model | 24hpir_brain | GLF | GO:0005515 |
|  | ENSDARG00000104387 | slc4a5b | full_model | 24hpir_intestine | GLC | GO:0008509 |
|  | ENSDARG00000079105 | mhc2dab | full_model | 24hpir_intestine | GLC | GO:0042613 |
|  | ENSDARG00000104635 | mhc2dgb | full_model | 24hpir_intestine | GLC |  |
|  | ENSDARG00000036045 | penkb | full_model | 24hpir_intestine | GLC | GO:0007218 |
|  | ENSDARG00000103716 | mhc2dga | full_model | 24hpir_intestine | GLC | GO:0042613 |
|  | ENSDARG00000016319 | c9 | full_model | 24hpir_intestine | GLF |  |
|  | ENSDARG00000068621 | nlrc11 | full_model | 24hpir_intestine | GLF | GO:0005515 |
|  | ENSDARG00000012694 | c3a.1 | full_model | 24hpir_intestine | GLF | GO:0004866 |
|  | ENSDARG00000055278 | cfb | full_model | 24hpir_intestine | GLF | GO:0006956 |
|  | ENSDARG00000102456 | cfhl4 | full_model | 24hpir_intestine | GLF |  |
|  | ENSDARG00000079105 | mhc2dab | full_model | 24hpir_kidney | GLC | GO:0042613 |
|  | ENSDARG00000091280 | si:ch211-66k16.27 | full_model | 24hpir_kidney | GLC | GO:0042981 |
|  | ENSDARG00000067848 | nmrk2 | full_model | 24hpir_kidney | GLC | GO:0005737 |
|  | ENSDARG00000038587 | LOC100004199 | full_model | 24hpir_kidney | GLC |  |
|  | ENSDARG00000037550 | cyp21a2 | full_model | 24hpir_kidney | GLC | GO:0016705 |
|  | ENSDARG00000068621 | nlrc11 | full_model | 24hpir_kidney | GLF | GO:0005515 |
|  | ENSDARG00000091847 | LOC110437747 | full_model | 24hpir_kidney | GLF | GO:0005515 |
|  | ENSDARG00000094210 | fthl31 | full_model | 24hpir_kidney | GLF | GO:0006879 |
|  | ENSDARG00000016771 | tfa | full_model | 24hpir_kidney | GLF |  |
|  | ENSDARG00000086569 | zgc:172051 | full_model | 24hpir_kidney | GLF |  |
|  | ENSDARG00000079105 | mhc2dab | full_model | 24hpir_caudal fin | GLC | GO:0042613 |
|  | ENSDARG00000042988 | slc24a2 | full_model | 24hpir_caudal fin | GLC |  |
|  | ENSDARG00000104635 | mhc2dgb | full_model | 24hpir_caudal fin | GLC |  |
|  | ENSDARG00000044074 | loxl2b | full_model | 24hpir_caudal fin | GLC | GO:0016641 |
|  | ENSDARG00000095082 | si:ch211-194p6.12 | full_model | 24hpir_caudal fin | GLC |  |
|  | ENSDARG00000068621 | nlrc11 | full_model | 24hpir_caudal fin | GLF | GO:0005515 |
|  | ENSDARG00000091847 | LOC110437747 | full_model | 24hpir_caudal fin | GLF | GO:0005515 |
|  | ENSDARG00000062788 | irg1l | full_model | 24hpir_caudal fin | GLF | GO:0016829 |
|  | ENSDARG00000116832 | NA | full_model | 24hpir_caudal fin | GLF | GO:0015074 |
|  | ENSDARG00000007769 | sult5a1 | full_model | 24hpir_caudal fin | GLF | GO:0016740 |
|  | ENSDARG00000079105 | mhc2dab | full_model | 24hpir_spleen | GLC | GO:0042613 |
|  | ENSDARG00000104635 | mhc2dgb | full_model | 24hpir_spleen | GLC |  |
|  | ENSDARG00000038587 | LOC100004199 | full_model | 24hpir_spleen | GLC |  |
|  | ENSDARG00000116524 | mhc2dhb | full_model | 24hpir_spleen | GLC | GO:0042613 |
|  | ENSDARG00000071024 | zgc:171679 | full_model | 24hpir_spleen | GLC | GO:0005515 |
|  | ENSDARG00000020084 | tg | full_model | 24hpir_spleen | GLF |  |
|  | ENSDARG00000068621 | nlrc11 | full_model | 24hpir_spleen | GLF | GO:0005515 |
|  | ENSDARG00000069296 | moxd1l | full_model | 24hpir_spleen | GLF | GO:0004500 |
|  | ENSDARG00000091847 | LOC110437747 | full_model | 24hpir_spleen | GLF | GO:0005515 |
|  | ENSDARG00000076043 | si:dkeyp-73d8.9 | full_model | 24hpir_spleen | GLF |  |
|  | ENSDARG00000015866 | apoa2 | germline | 24hpir_tests | GLC |  |
|  | ENSDARG00000075261 | timp2b | germline | 24hpir_tests | GLC | GO:0008191 |
|  | ENSDARG00000003533 | col8a1b | germline | 24hpir_tests | GLC | GO:0005515 |
|  | ENSDARG00000069282 | bbc3 | germline | 24hpir_tests | GLC | GO:0090200 |
|  | ENSDARG00000101324 | apoa1b | germline | 24hpir_tests | GLC | GO:0042157 |

#### Table S6. Mixed-effects modelling of somatic gene structural properties. Results from LMMs (gene expression, gene length) and GLMMs (exonic proportion) fitted to the full somatic dataset. Fixed and random effects are reported.

**A) Gene expression (logCPM)**

| **Effect** | **χ²** | **df** | **p** |
| --- | --- | --- | --- |
| Strain | 63 | 1 | < 0.0001 |
| Contrast | 2,412 | 17 | < 0.0001 |
| Gene length (covariate) | 770 | 1 | < 0.0001 |
| Chromatin openness | 11 | 1 | < 0.001 |
| Strain × Contrast | 2,252 | 17 | < 0.0001 |
| Random effects: chromosome variance 0.08 (SD 0.28); strand variance 0.001 (SD 0.03) |  |  |  |

**B) Gene length (bp)**

| **Effect** | **χ²** | **df** | **p** |
| --- | --- | --- | --- |
| Strain | 34 | 1 | < 0.0001 |
| Contrast | 717 | 17 | < 0.0001 |
| Chromatin openness | 7,052 | 1 | < 0.0001 |
| Strain × Contrast | 1,538 | 17 | < 0.0001 |
| Random effects: chromosome variance 0.007 (SD 0.08); strand variance 0.0002 (SD 0.01) |  |  |  |

**C) Exonic proportion (%)**

| **Effect** | **χ²** | **df** | **p** |
| --- | --- | --- | --- |
| Strain | 18,731 | 1 | < 0.0001 |
| Contrast | 146,372 | 17 | < 0.0001 |
| Gene length (covariate) | 124,737,573 | 1 | < 0.0001 |
| Chromatin openness | 321,416 | 1 | < 0.0001 |
| Strain × Contrast | 301,334 | 17 | < 0.0001 |
| Random effects: chromosome variance 0.005 (SD 0.07); strand variance 0.0001 (SD 0.10) |  |  |  |

#### Table S7. Pairwise comparisons of estimated marginal means. Pairwise contrasts from LMMs (gene expression, gene length) and GLMMs (exonic proportion). P-values adjusted by Tukey HSD.

**A) Gene expression (logCPM)**

| **Contrast** | **Estimate** | **SE** | **df** | **z-ratio** | **p-value** |
| --- | --- | --- | --- | --- | --- |
| GLC C1 - GLF C1 | -0.492 | 0.062 | Inf | -7.932 | <.0001 |
| GLC C2.3 - GLF C2.3 | 0.725 | 0.05392563 | Inf | 13.442 | <.0001 |
| GLC C2.24 - GLF C2.24 | -0.711 | 0.0553 | Inf | -12.874 | <.0001 |
| GLC C1 - GLC C2.3 | -0.572 | 0.052 | Inf | -11.097 | <.0001 |
| GLC C1 - GLC C2.24 | 0.409 | 0.051 | Inf | 8.009 | <.0001 |
| GLC C2.24 - GLC C2.3 | -0.981 | 0.051 | Inf | -19.167 | <.0001 |
| GLF C1 - GLF C2.3 | 0.645 | 0.064 | Inf | 10.077 | <.0001 |
| GLF C1 - GLF C2.24 | 0.190 | 0.065 | Inf | 2.898 | <.0001 |
| GLF C2.24 - GLF C2.3 | 0.455 | 0.058 | Inf | 7.882 | <.0001 |
| GLC C3.br - GLF C3.br | -0.249 | 0.073 | Inf | -3.428 | <.0001 |
| GLC C3.int - GLF C3.int | -0.451 | 0.053 | Inf | -8.564 | <.0001 |
| GLC C3.kid - GLF C3.kid | -0.416 | 0.051 | Inf | -8.133 | <.0001 |
| GLC C3.ski - GLF C3.ski | 0.108 | 0.051 | Inf | 2.123 | <.0001 |
| GLC C3.spl - GLF C3.spl | -0.533 | 0.051 | Inf | -10.465 | <.0001 |
| GLC C4.br.3 - GLF C4.br.3 | 1.387 | 0.053 | Inf | 26.314 | <.0001 |
| GLC C4.int.3 - GLF C4.int.3 | 0.2821 | 0.0491 | Inf | 5.7407 | <.0001 |
| GLC C4.kid.3 - GLF C4.kid.3 | -0.048 | 0.049 | Inf | -0.979 | <.0001 |
| GLC C4.ski.3 - GLF C4.ski.3 | 0.765 | 0.050 | Inf | 15.238 | <.0001 |
| GLC C4.spl.3 - GLF C4.spl.3 | 0.9058 | 0.0459 | Inf | 19.7469 | <.0001 |
| GLC C5.br.24 - GLF C5.br.24 | -0.258 | 0.071 | Inf | -3.611 | <.0001 |
| GLC C5.int.24 - GLF C5.int.24 | -0.525 | 0.047 | Inf | -11.236 | <.0001 |
| GLC C5.kid.24 - GLF C5.kid.24 | -0.367 | 0.046 | Inf | -8.000 | <.0001 |
| GLC C5.ski.24 - GLF C5.ski.24 | -0.129 | 0.046 | Inf | -2.786 | <.0001 |
| GLC C5.spl.24 - GLF C5.spl.24 | -0.307 | 0.054 | Inf | -5.681 | <.0001 |

**B) Gene length**

| **Contrast** | **Estimate** | **SE** | **df** | **z-ratio** | **p-value** |
| --- | --- | --- | --- | --- | --- |
| GLC C1 - GLF C1 | 0.1793 | 0.0309 | Inf | 5.8037 | <.0001 |
| GLC C2.3 - GLF C2.3 | -0.3542 | 0.0268 | Inf | -13.1951 | <.0001 |
| GLC C2.24 - GLF C2.24 | 0.1444 | 0.0275 | Inf | 5.2462 | <.0001 |
| GLC C1 - GLC C2.3 | 0.2508 | 0.0256 | Inf | 9.7795 | <.0001 |
| GLC C1 - GLC C2.24 | 0.0307 | 0.0254 | Inf | 1.2079 | <.0001 |
| GLC C2.24 - GLC C2.3 | 0.2201 | 0.0255 | Inf | 8.6381 | <.0001 |
| GLF C1 - GLF C2.3 | -0.2826 | 0.0319 | Inf | -8.8646 | <.0001 |
| GLF C1 - GLF C2.24 | -0.0042 | 0.0326 | Inf | -0.1274 | <.0001 |
| GLF C2.24 - GLF C2.3 | -0.2784 | 0.0288 | Inf | -9.6776 | <.0001 |
| GLC C3.br - GLF C3.br | 0.3898 | 0.0361 | Inf | 10.7859 | <.0001 |
| GLC C3.int - GLF C3.int | 0.1611 | 0.0262 | Inf | 6.1448 | <.0001 |
| GLC C3.kid - GLF C3.kid | 0.0510 | 0.0255 | Inf | 1.9999 | <.0001 |
| GLC C3.ski - GLF C3.ski | 0.0344 | 0.0253 | Inf | 1.3622 | <.0001 |
| GLC C3.spl - GLF C3.spl | 0.0758 | 0.0253 | Inf | 2.9904 | <.0001 |
| GLC C4.br.3 - GLF C4.br.3 | -0.5949 | 0.0262 | Inf | -22.6935 | <.0001 |
| GLC C4.int.3 - GLF C4.int.3 | -0.0864 | 0.0245 | Inf | -3.5301 | <.0001 |
| GLC C4.kid.3 - GLF C4.kid.3 | -0.0889 | 0.0246 | Inf | -3.6125 | <.0001 |
| GLC C4.ski.3 - GLF C4.ski.3 | -0.3537 | 0.0250 | Inf | -14.1572 | <.0001 |
| GLC C4.spl.3 - GLF C4.spl.3 | -0.3856 | 0.0228 | Inf | -16.8984 | <.0001 |
| GLC C5.br.24 - GLF C5.br.24 | 0.2516 | 0.0356 | Inf | 7.0681 | <.0001 |
| GLC C5.int.24 - GLF C5.int.24 | 0.1738 | 0.0233 | Inf | 7.4723 | <.0001 |
| GLC C5.kid.24 - GLF C5.kid.24 | 0.0165 | 0.0228 | Inf | 0.7247 | <.0001 |
| GLC C5.ski.24 - GLF C5.ski.24 | 0.1368 | 0.0230 | Inf | 5.9434 | <.0001 |
| GLC C5.spl.24 - GLF C5.spl.24 | 0.0195 | 0.0269 | Inf | 0.7238 | <.0001 |

**A) Proportion of exonic sequence**

| **Contrast** | **Estimate** | **SE** | **df** | **z-ratio** | **p-value** |
| --- | --- | --- | --- | --- | --- |
| GLC C1 - GLF C1 | -0.0732 | 0.0005 | Inf | -136.8623 | <.0001 |
| GLC C2.3 - GLF C2.3 | 0.0358 | 0.0005 | Inf | 77.2684 | <.0001 |
| GLC C2.24 - GLF C2.24 | -0.0537 | 0.0005 | Inf | -111.2968 | <.0001 |
| GLC C1 - GLC C2.3 | -0.0499 | 0.0004 | Inf | -111.3901 | <.0001 |
| GLC C1 - GLC C2.24 | 0.0077 | 0.0004 | Inf | 17.6583 | <.0001 |
| GLF C1 - GLF C2.3 | 0.0591 | 0.0005 | Inf | 107.8751 | <.0001 |
| GLF C1 - GLF C2.24 | 0.0272 | 0.0006 | Inf | 47.3808 | <.0001 |
| GLF C2.24 - GLF C2.3 | 0.0319 | 0.0005 | Inf | 64.2225 | <.0001 |
| GLC C3.br - GLF C3.br | -0.1395 | 0.0007 | Inf | -210.0547 | <.0001 |
| GLC C3.int - GLF C3.int | -0.0271 | 0.0005 | Inf | -57.3460 | <.0001 |
| GLC C3.kid - GLF C3.kid | -0.0704 | 0.0004 | Inf | -158.4571 | <.0001 |
| GLC C3.ski - GLF C3.ski | -0.0305 | 0.0004 | Inf | -67.8595 | <.0001 |
| GLC C3.spl - GLF C3.spl | -0.0809 | 0.0005 | Inf | -178.1585 | <.0001 |
| GLC C4.br.3 - GLF C4.br.3 | 0.0932 | 0.0005 | Inf | 193.8331 | <.0001 |
| GLC C4.int.3 - GLF C4.int.3 | 0.0443 | 0.0004 | Inf | 101.3887 | <.0001 |
| GLC C4.kid.3 - GLF C4.kid.3 | 0.0037 | 0.0004 | Inf | 8.4026 | <.0001 |
| GLC C4.ski.3 - GLF C4.ski.3 | 0.0419 | 0.0004 | Inf | 94.7956 | <.0001 |
| GLC C4.spl.3 - GLF C4.spl.3 | 0.0941 | 0.0004 | Inf | 232.0648 | <.0001 |
| GLC C5.br.24 - GLF C5.br.24 | -0.0599 | 0.0006 | Inf | -92.7293 | <.0001 |
| GLC C5.int.24 - GLF C5.int.24 | -0.0443 | 0.0004 | Inf | -106.2674 | <.0001 |
| GLC C5.kid.24 - GLF C5.kid.24 | -0.0533 | 0.0004 | Inf | -130.1877 | <.0001 |
| GLC C5.ski.24 - GLF C5.ski.24 | 0.0219 | 0.0004 | Inf | 53.4500 | <.0001 |
| GLC C5.ski.24 - GLF C5.spl.24 | -0.0247 | 0.0004 | Inf | -56.3571 | <.0001 |

#### Table S8. Mixed-effects modelling of germline gene structural properties. LMMs fitted to germline (testes) data only. Fixed effects of irradiation, gene length, and chromatin openness; random effect of chromosome.

**A) Gene expression (logCPM)**

| **Effect** | **χ²** | **df** | **p** |
| --- | --- | --- | --- |
| Irradiation | 132 | 1 | < 0.0001 |
| Gene length (covariate) | 15 | 1 | < 0.0001 |
| Chromatin openness | 20 | 1 | < 0.0001 |
| Random effects: chromosome variance 0.08 (SD 0.30) |  |  |  |

**B) Gene length (bp)**

| **Effect** | **χ²** | **df** | **p** |
| --- | --- | --- | --- |
| Irradiation | 498 | 1 | < 0.0001 |
| Chromatin openness | 1,277 | 1 | < 0.0001 |
| Random effects: chromosome variance 0.008 (SD 0.09) |  |  |  |

**C) Exonic proportion (%)**

| **Effect** | **χ²** | **df** | **p** |
| --- | --- | --- | --- |
| Irradiation | 25,951 | 1 | < 0.0001 |
| Gene length (covariate) | 23,358,721 | 1 | < 0.0001 |
| Chromatin openness | 36,931 | 1 | < 0.0001 |
| Random effects: chromosome variance 0.005 (SD 0.07); strand variance 0.00008 (SD 0.009) |  |  |  |

#### Table S9. Gene index for the ageing GO term alluvial plot (Figure 4). All genes contributing to alluvial flows shown in Figure 4 are listed by timepoint (non-irradiated, 3 hpir, 24 hpir) to enable identification of individual genes underlying each pathway–strain–timepoint interaction. Full gene list available at the project GitHub repository.

| Gene Index | nonir | 3hpir | 24hpir | Gene Index | nonir | 3hpir | 24hpir | Gene Index | nonir | 3hpir | 24hpir |
| --- | --- | --- | --- | --- | --- | --- | --- | --- | --- | --- | --- |
| **1** | abat | abat | abl1 | **35** | foxm1 | gprc5c | cyt1 | **63** | nrg1 | pitx3 | myl6 |
| **2** | abl1 | agt | acta2 | **36** | foxo3b | hamp | dag1 | **64** | park7 | ppp3ca | myo5b |
| **3** | ace | amh | agt | **37** | fxn | il23r | ddc | **65** | pawr | prelp | myo5c |
| **4** | aco1 | arg2 | akap9 | **38** | gas6 | irak1 | dnmbp | **66** | pck1 | rad54b | nedd4l |
| **5** | aco2 | atr | appl2 | **39** | gprc5c | jund | eef1e1 | **67** | pibf1 | rad54l | nek6 |
| **6** | adsl | aurkb | atr | **40** | ikbkb | kif14 | eif2b3 | **68** | pik3ca | rgcc | nono |
| **7** | agap2 | cab39 | b2m | **41** | il6r | kmo | endog | **69** | ppp2r3c | ripk3 | pck1 |
| **8** | agt | calca | bcar1 | **42** | inpp5d | lmna | fbxo4 | **70** | pqbp1 | romo1 | pde4d |
| **9** | ak4 | calr | blnk | **43** | ireb2 | lrrk2 | fcho1 | **71** | prkdc | rplp1 | pik3cd |
| **10** | atp7a | casp9 | brca2 | **44** | itgb1bp1 | map2k1 | fgf2 | **72** | pycard | rpn2 | pkp2 |
| **11** | atp8a2 | cd68 | c1qa | **45** | kat6a | map3k3 | flna | **73** | rgcc | rwdd1 | pla2r1 |
| **12** | atr | cdk2 | c1qc | **46** | malt1 | map3k5 | foxo3b | **74** | ripk3 | scap | polb |
| **13** | bnip3 | chek2 | c4b | **47** | map3k3 | mapkapk5 | gata3 | **75** | rptor | sec63 | pqbp1 |
| **14** | c1qa | cisd2 | c5 | **48** | mbl2 | mbd2 | gbp1 | **76** | sdhaf2 | serpine1 | prkd2 |
| **15** | calca | cited2 | c8a | **49** | mob3c | mif | gbp2 | **77** | sdhc | serpinf1 | ptprc |
| **16** | calr | cln8 | c8g | **50** | msh6 | mmp2 | gclc | **78** | shmt2 | slc1a1 | pycard |
| **17** | canx | cnp | cacna1g | **51** | nckap1l | mob3c | gpd1l | **79** | sirt3 | slc30a10 | rbl1 |
| **18** | card14 | col4a2 | calca | **52** | ndnl2 | ndnl2 | hck | **80** | slc1a1 | smc5 | rc3h2 |
| **19** | casp9 | comp | calr | **53** | ndufa2 | nsmce2 | irak1 | **81** | slc25a33 | sod1 | rela |
| **20** | cat | cops8 | cat | **54** | ndufa6 | nudt1 | itk | **82** | src | srebf1 | serping1 |
| **21** | chchd10 | ctc1 | cblb | **55** | ndufa7 | p2ry1 | klhl6 | **83** | stoml2 | srr | skap1 |
| **22** | cisd2 | ctnna1 | ccdc88c | **56** | ndufaf1 | park7 | lck | **84** | suclg2 | stk11 | slc12a2 |
| **23** | comp | ctsc | cd276 | **57** | ndufb2 | pawr | malt1 | **85** | tbk1 | stradb | src |
| **24** | cox15 | daxx | cd68 | **58** | ndufb3 | pck1 | mapk1 | **86** | tlr1 | tal1 | srebf1 |
| **25** | cox4i2 | dbf4b | cdk2 | **59** | ndufb6 | pdx1 | mbl2 | **87** | tlr3 | tert | srr |
| **26** | cox6a1 | ddc | cdk6 | **60** | ndufc1 | phb | mybpc3 | **88** | tomm70a | tfcp2l1 | susd4 |
| **27** | cox6a2 | dnmbp | cfb | **61** | ndufs5 | pibf1 | myh10 | **89** | tpx2 | tgfb3 | thy1 |
| **28** | cyc1 | edn1 | cfi | **62** | ndufv3 | pink1 | myh6 | **90** | traf6 | tlr1 | tnfrsf21 |
| **29** | dag1 | ercc1 | clu | **63** | nrg1 | pitx3 | myl6 | **91** | trpv4 | tlr3 | tomm70a |
| **30** | edn1 | fbxo4 | cmklr1 | **64** | park7 | ppp3ca | myo5b | **92** | uqcrc1 | tp53 | tp53 |
| **31** | emp2 | fgf2 | cmtm3 | **65** | pawr | prelp | myo5c | **93** | uqcrfs1 | tpra1 | tyrobp |
| **32** | endog | foxm1 | colec11 | **66** | pck1 | rad54b | nedd4l | **94** | wnt11 | ucp2 | ucp2 |
| **33** | ercc1 | foxo3b | csk | **67** | pibf1 | rad54l | nek6 | **95** | znfx1 | ulk3 | ulk3 |
| **34** | fh | gclc | ctc1 | **68** | pik3ca | rgcc | nono | **96** |  | vash1 | vash1 |

### Supplementary Figures

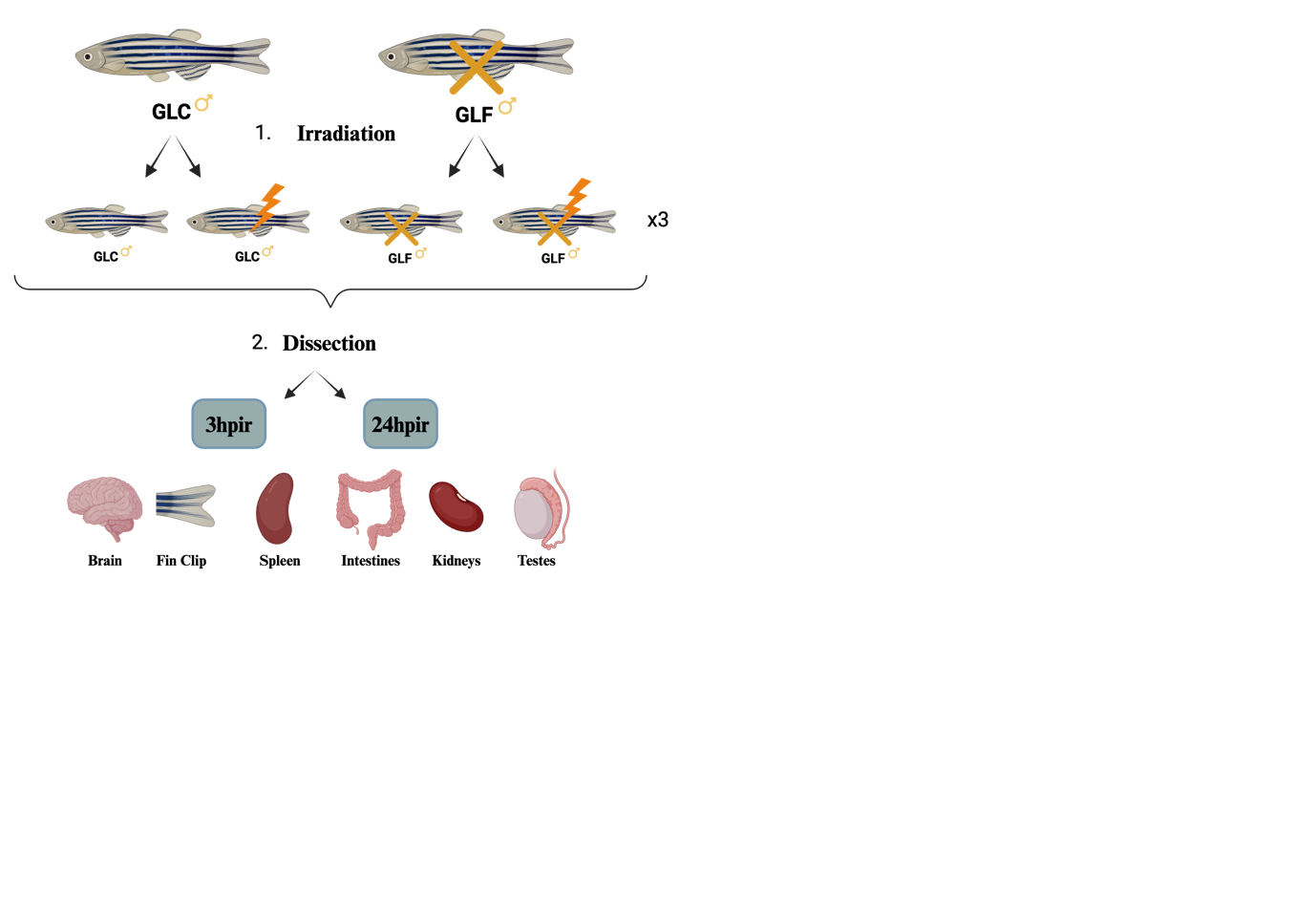

**Figure S1.** Experimental design. Schematic illustrating the full-factorial sampling design. Three males per treatment group (GLC or GLF) × irradiation condition (non-irradiated or irradiated) × timepoint (3 hpir or 24 hpir) were sampled, yielding 24 males and 126 tissue samples in total. Five somatic tissues (brain, intestine, kidney, spleen, caudal fin) were collected from all fish; testes were collected from GLC fish only. Tissues were dissected on ice and flash-frozen in liquid nitrogen immediately after collection.

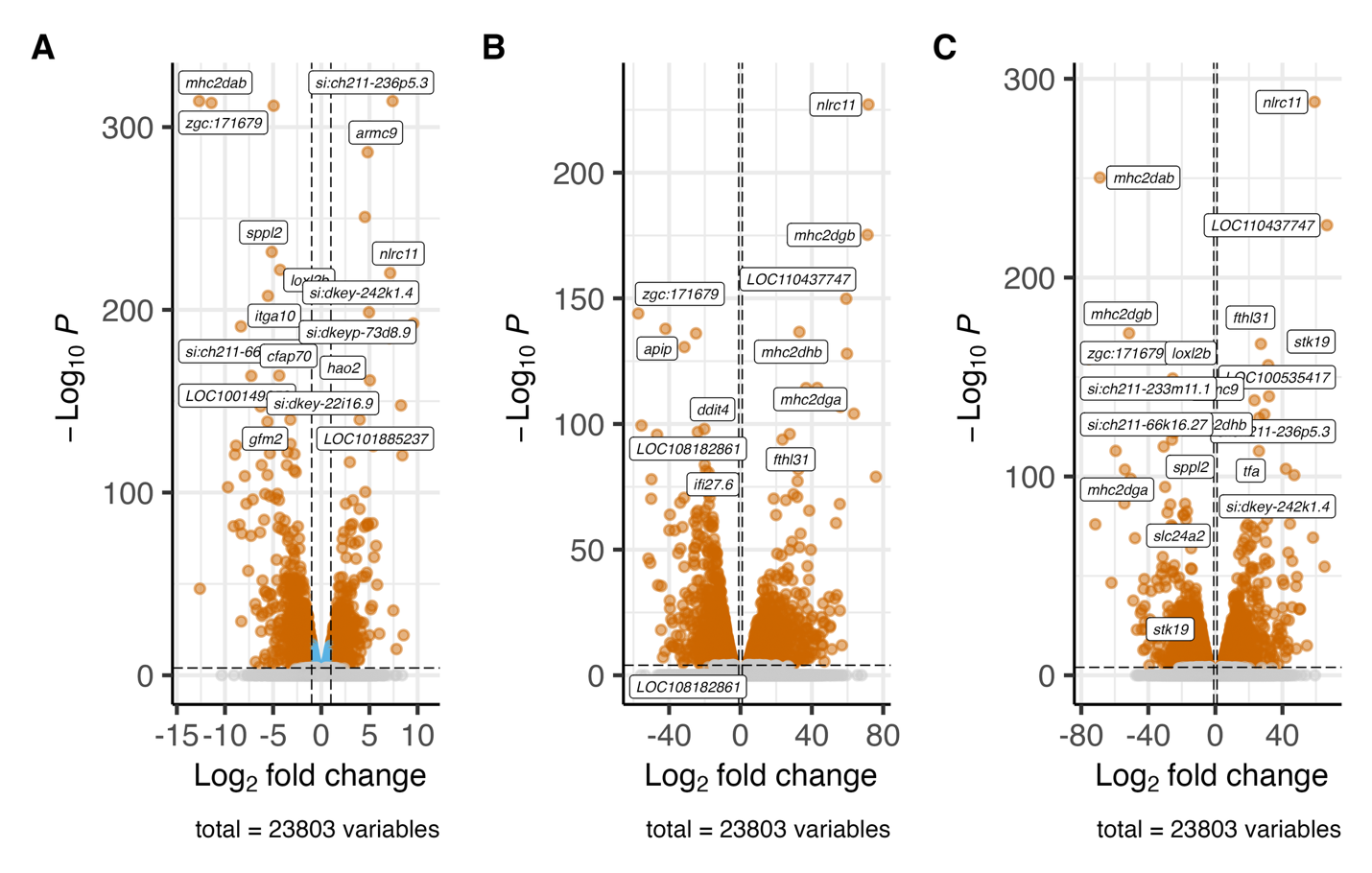

**Figure S2.** Volcano plots of somatic differential expression for the three soma-level contrasts. Each point represents one gene. Genes with positive log₂ fold change (logFC) are expressed at higher levels in GLF; those with negative logFC are expressed at higher levels in GLC. Non-significant genes (FDR ≥ 0.05) are shown in grey. Horizontal and vertical dashed lines mark the significance threshold (−log₁₀(p-value) = 1.3) and fold-change thresholds (log₂FC = ±0.6), respectively. **(A)** Contrast C1 (non-irradiated soma); **(B)** contrast C2.3 (3 hpir soma); **(C)** contrast C2.24 (24 hpir soma).

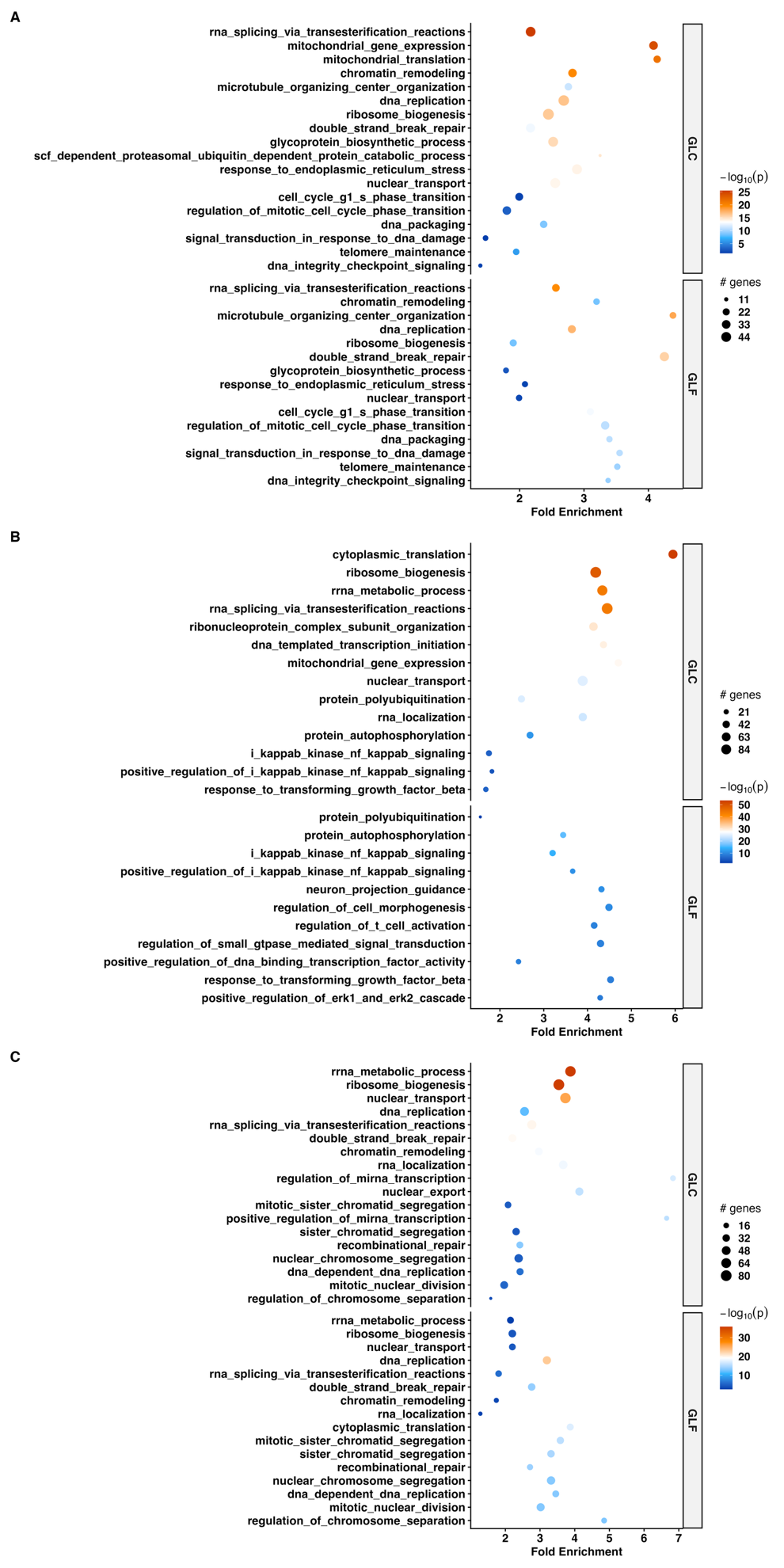

**Figure S3.** GO term enrichment across irradiation timepoints reveals strain-dependent shifts in genome maintenance and metabolic programmes. Bubble plots of significantly enriched GO terms at **(A)** non-irradiated, **(B)** 3 hpir, and **(C)** 24 hpir timepoints, stratified by strain. Bubble colour indicates statistical significance (−log₁₀ *p-*value; orange = high, blue = low); bubble size reflects the number of associated genes. Only terms with *p_adj_* < 0.1 are displayed.

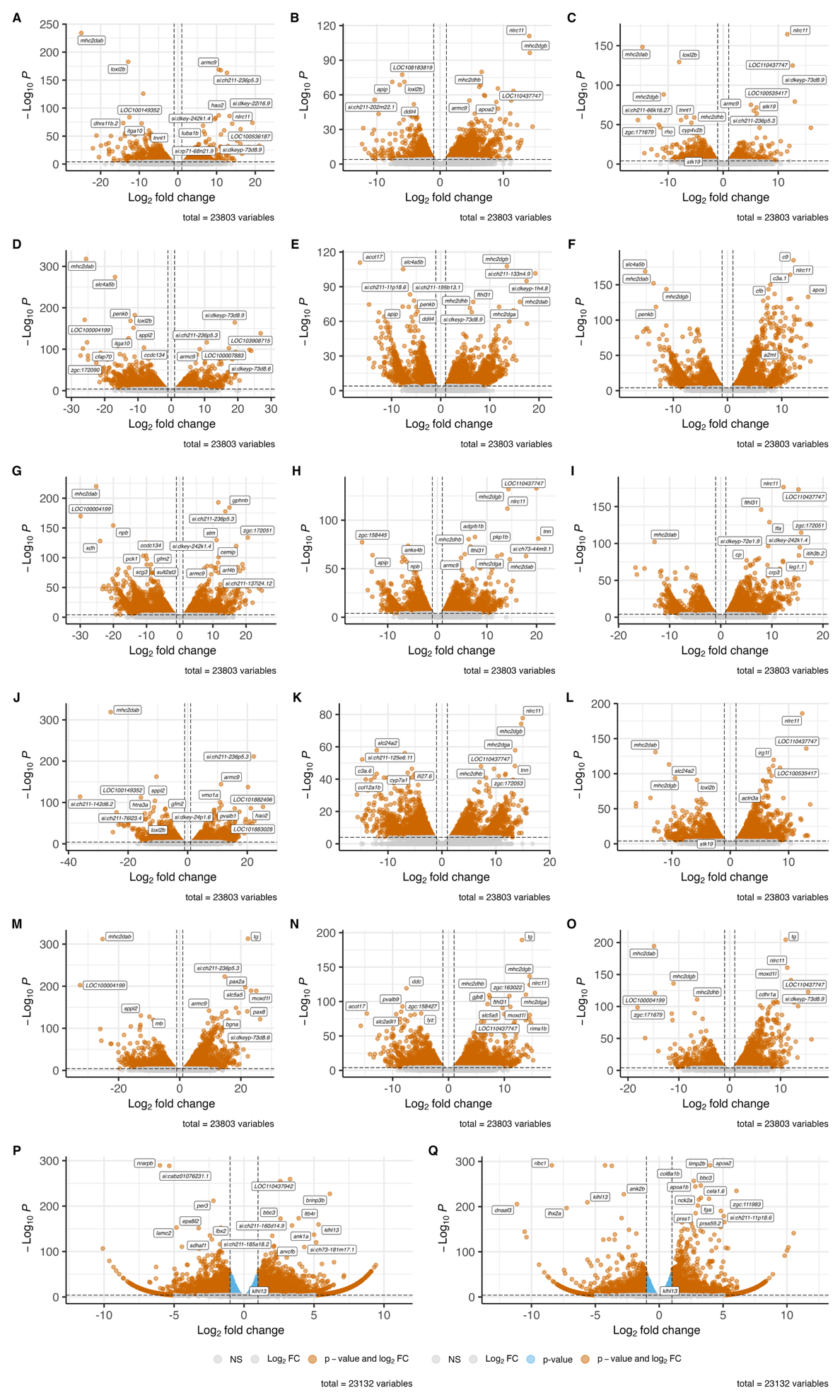

**Figure S4.** Volcano plots of differential expression across all tissues and timepoints. Format as described for Supplementary Figure S2. **(A–C)** Brain at non-irradiated, 3 hpir, and 24 hpir; **(D–F)** intestine; **(G–I)** kidney; (J–L) caudal fin; **(M–O)** spleen**. (P)** Contrast C6.3 (non-irradiated GLC testes vs 3 hpir GLC testes); **(Q)** contrast C6.24 (non-irradiated GLC testes vs 24 hpir GLC testes).

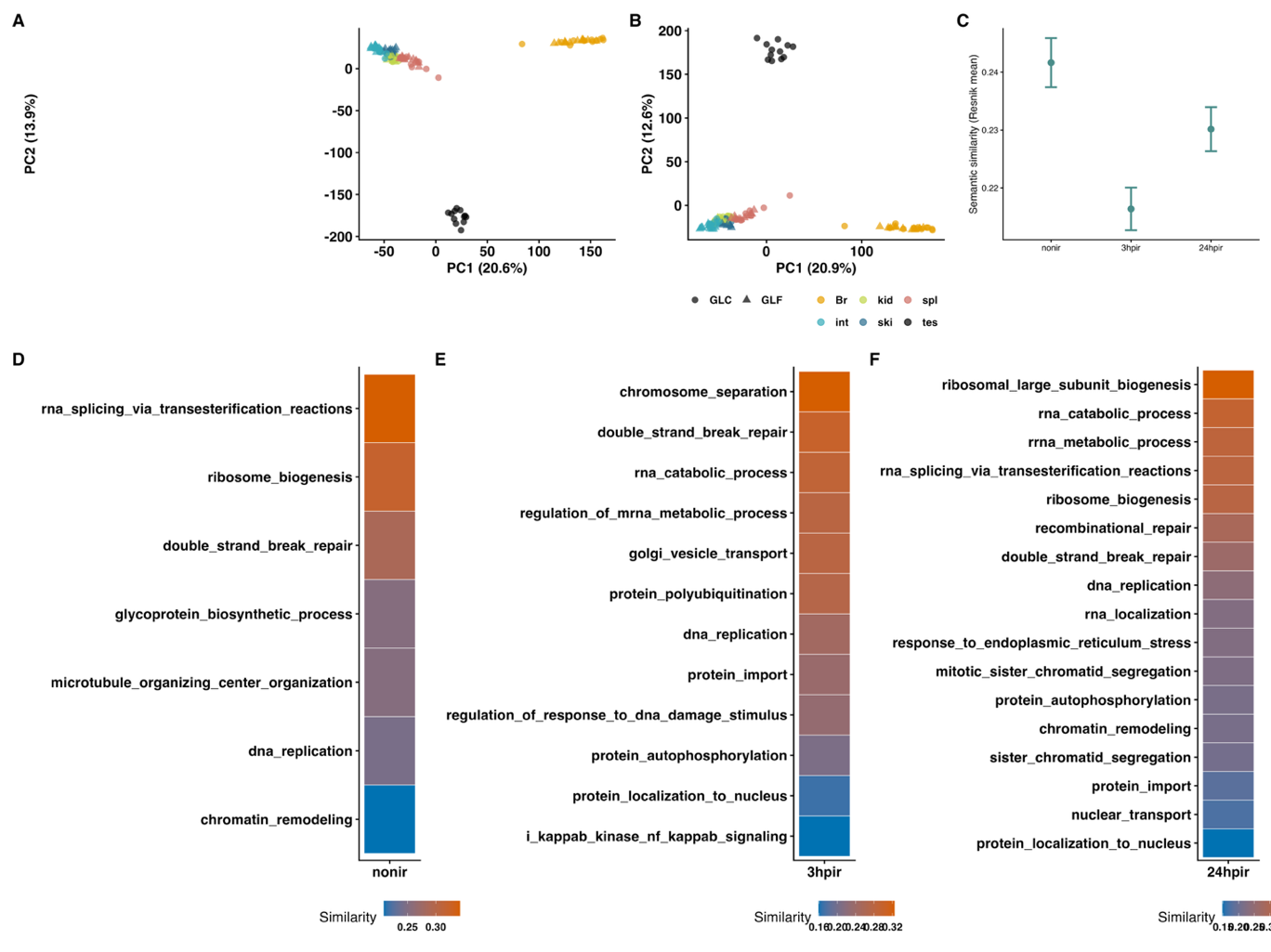

**Figure S5.** PCA and semantic similarity analysis of differentially expressed genes and transposable elements. **(A)** PCA of DE genes across all samples, with strain (GLC/GLF) represented by shape and tissue by colour. **(B)** PCA of DE TEs across the same samples. **(C)** Mean Resnik semantic similarity between GLC and GLF gene sets across timepoints; lower values indicate lower functional overlap. Resnik similarity heatmaps for the 100 most statistically enriched GO terms per condition: **(D)** non-irradiated, **(E)** 3 hpir, **(F)** 24 hpir. Colour intensity reflects semantic similarity between GLC and GLF gene sets; GO terms are ordered by increasing similarity within each condition.

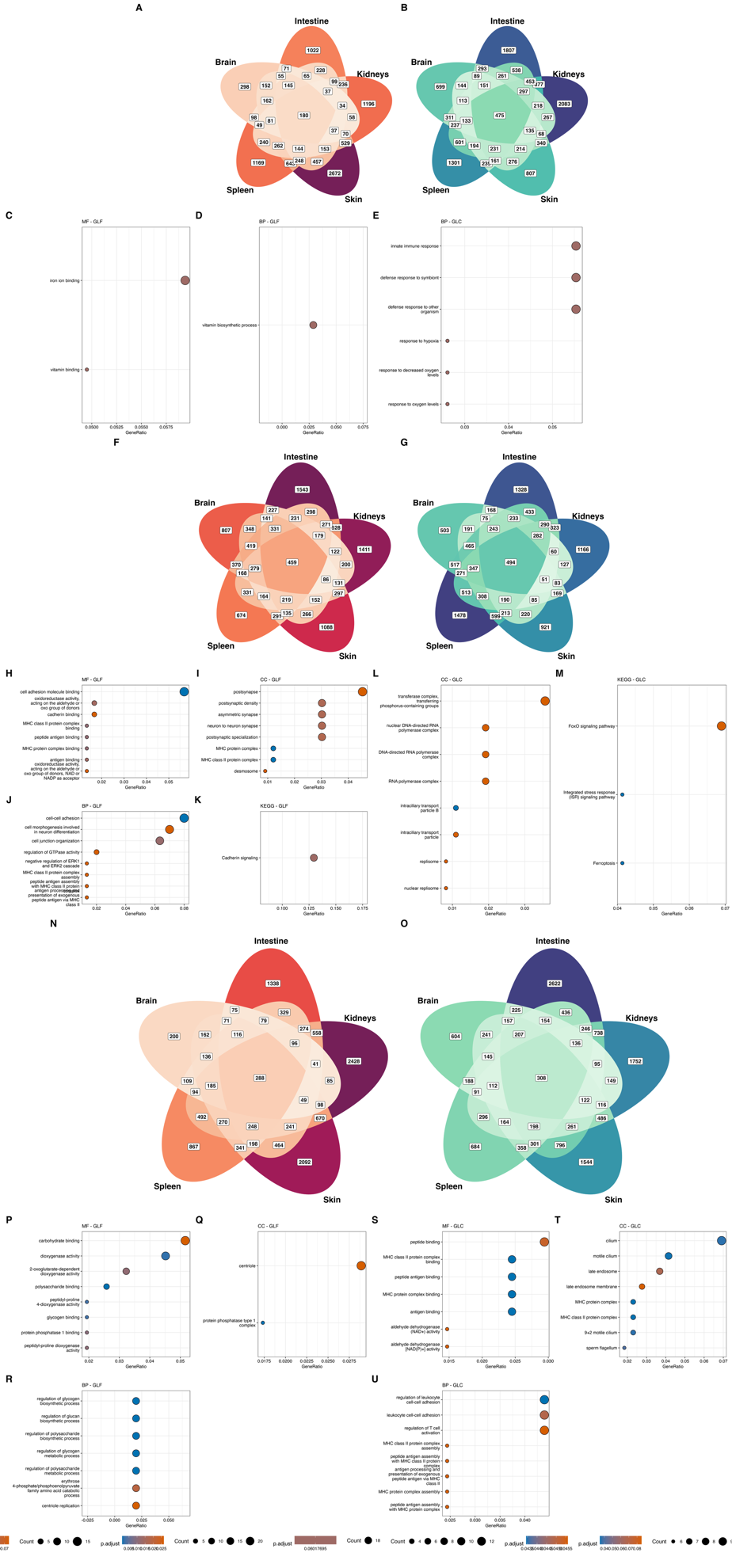

**Figure S6.** Functional enrichment of shared DE genes in somatic tissues across conditions. Venn diagrams of DE genes upregulated in GLF **(A)** and GLC **(B)** non-irradiated soma. Functional enrichment of 180 overlapping genes in GLF non-irradiated soma: **(C)** Molecular Function, **(D)** Biological Process, **(E)** GLC overlapping genes (n = 475) by Biological Process. Analogous enrichments for 3 hpir: GLF overlapping genes by **(H)** Molecular Function, **(I)** Cellular Component, **(J)** Biological Process; GLC by **(K)** Cellular Component, **(L)** KEGG. Enrichments for 24 hpir GLF with n = 288 genes **(M–Q**) and GLC with n = 308 genes (**R–T)**. Only terms with *p_adj_* < 0.1 are displayed.

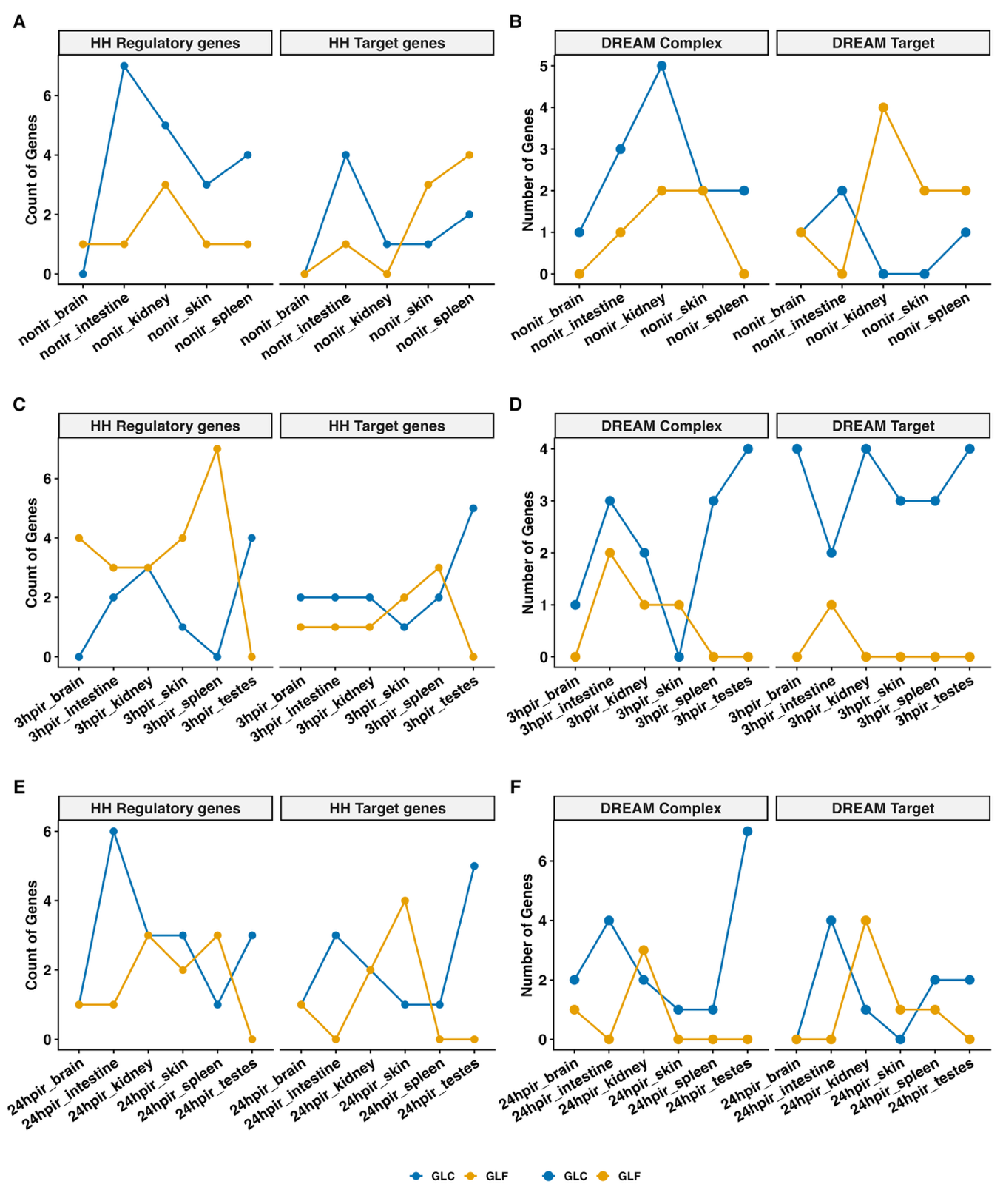

**Figure S7.** Tissue-level resolution of DREAM complex and Hedgehog pathway gene expression dynamics. Line plots showing DE gene counts (FDR < 0.05) per somatic tissue across timepoints for Hedgehog pathway components/targets (A, C, E) and DREAM complex components/targets **(B, D, F)**, at non-irradiated **(A–B)**, 3 hpir **(C–D)**, and 24 hpir **(E–F)** timepoints. Lines and points represent DE gene dynamics per tissue, coloured by strain.

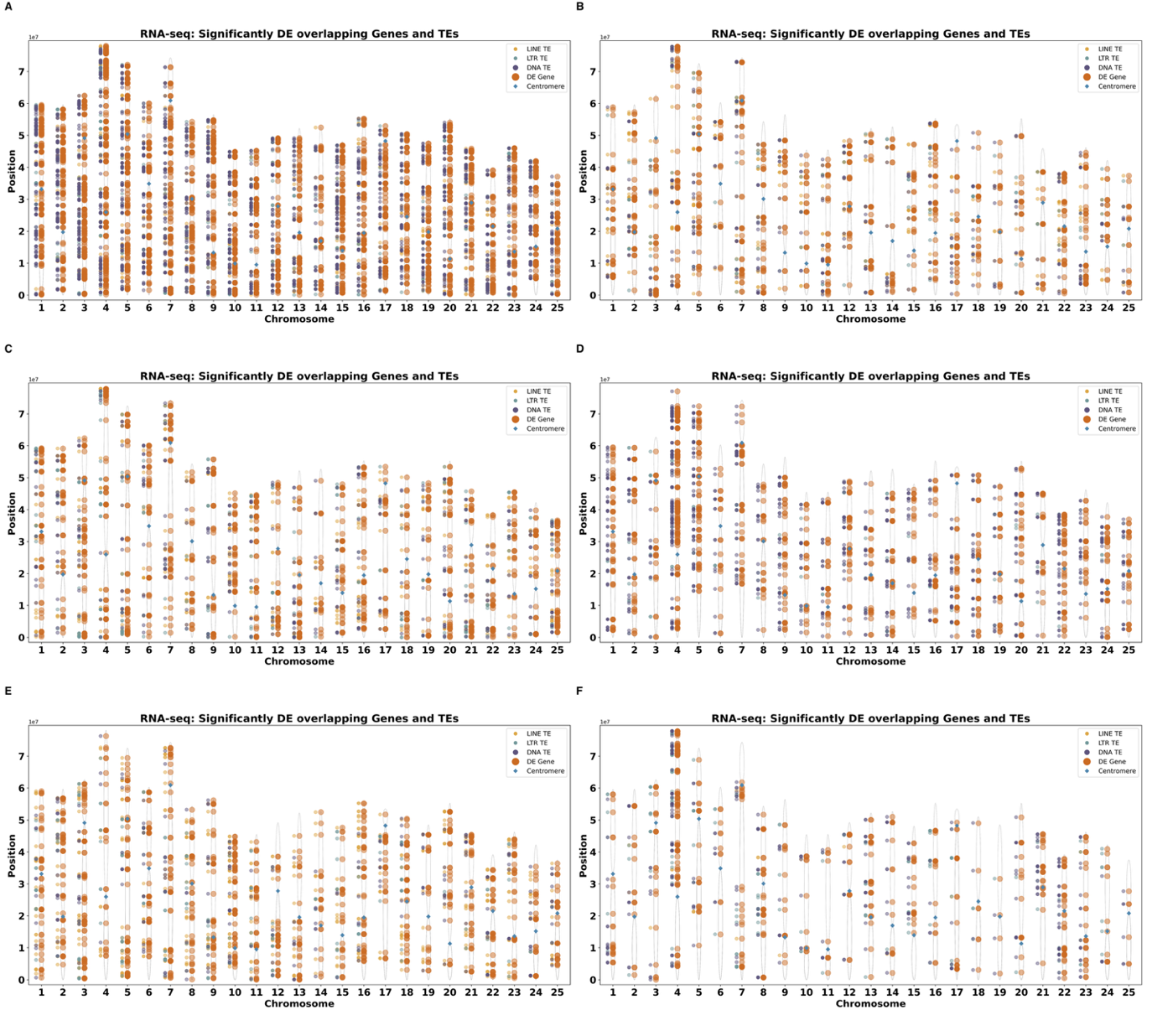

**Figure S8.** Chromosome 4 as a hotspot of coordinated DE gene–TE activity following irradiation in GLF fish. Ideogram-style plots for each strain × timepoint combination**: (A)** GLC non-irradiated, **(B)** GLF non-irradiated, **(C)** GLC 3 hpir, **(D)** GLF 3 hpir, **(E)** GLC 24 hpir, (F) GLF 24 hpir. DE genes are shown in orange; overlapping DE TEs are colour-coded by class: LINE (yellow), LTR (blue), DNA (purple). Each point corresponds to the genomic start position of a gene or TE, plotted against chromosome identity. TE positions are laterally offset for clarity. Centromere locations are marked with blue diamonds; grey ellipses approximate chromosome arms.

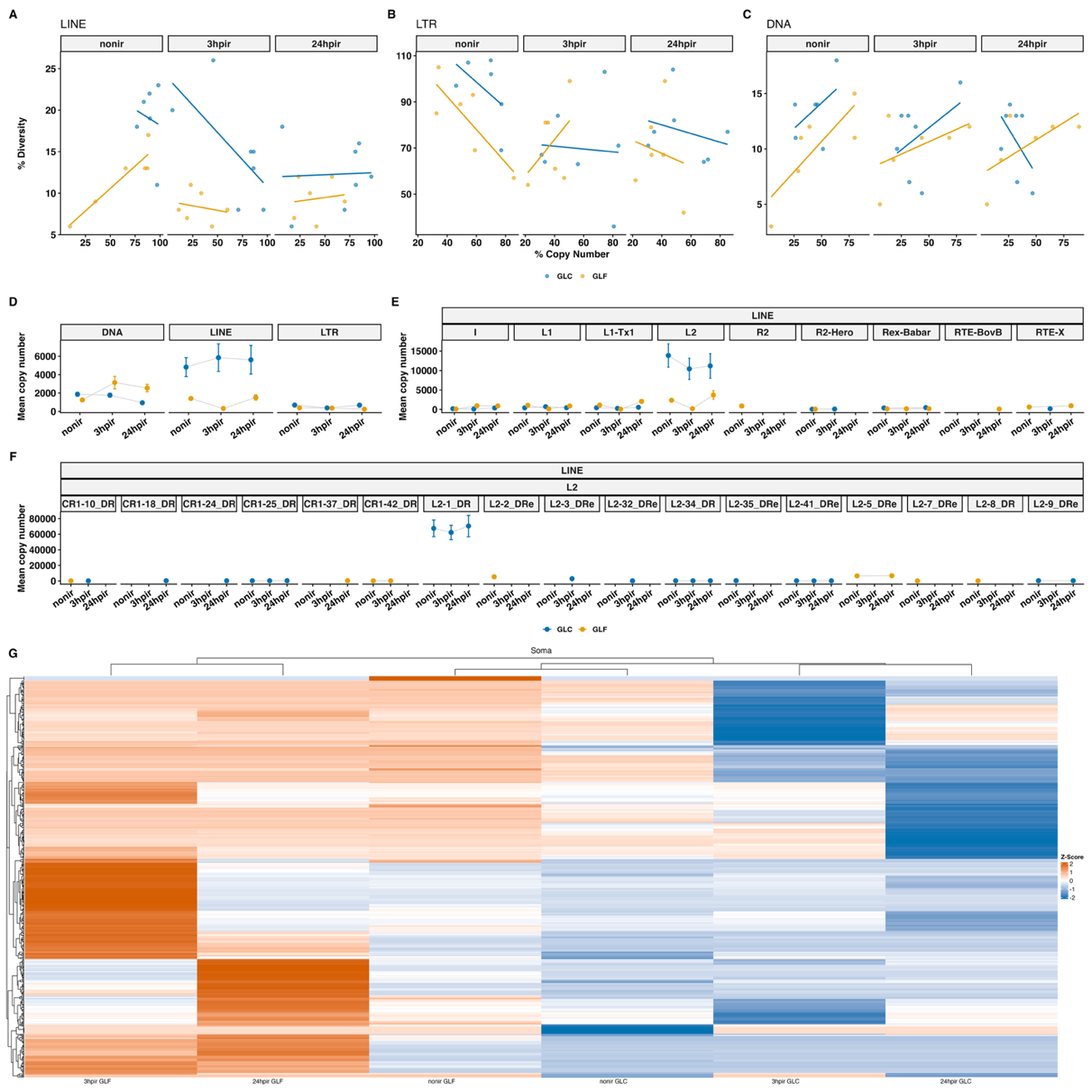

**Figure S9.** Strain-specific divergence in transposable element copy number, diversity, and expression dynamics. Scatterplots of the percentage of total copy number expressed (x-axis) vs number of significantly expressed repeats (y-axis) for LINE **(A)**, LTR **(B)**, and DNA **(C)** transposons. Each point represents a strain × timepoint contrast; linear models are shown per panel. **(D)** Mean copy number (± SE) of significantly expressed LINE, LTR, and DNA classes across timepoints, coloured by strain. **(E)** LINE subfamily copy number trajectories across time. **(F)** L2 subfamily copy number dynamics. **(G)** Heatmap of scaled logFC (Z-score) for DE TE repeats (rows) across somatic tissues (columns) for non-irradiated vs 3 hpir comparisons. Row Z-scores are normalised per TE repeat. Both rows and columns are hierarchically clustered. Orange indicates relative upregulation; blue indicates downregulation.

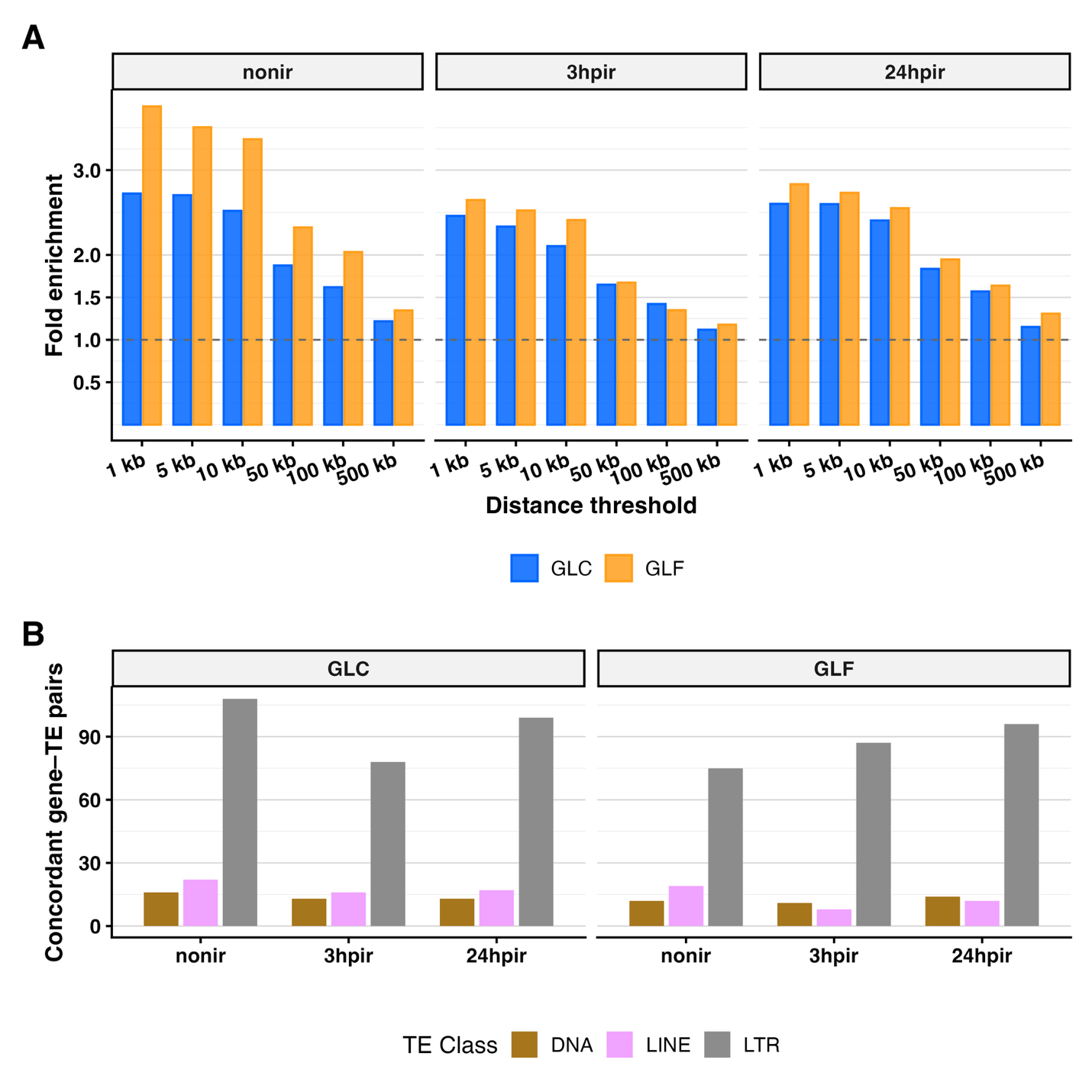

**Figure S10.** DE genes are enriched in proximity to DE transposable element loci and show concordant expression with LTR retrotransposons. **(A)** Fold enrichment of DE genes relative to all expressed genes within increasing genomic distance thresholds (1, 5, 10, 50, 100, 500 kb) of the nearest DE TE locus, across soma timepoints (non-irradiated, 3 hpir, 24 hpir) and stratified by strain (GLC, GLF). For each DE TE family, the single locus closest to any DE gene was retained. A dashed line at fold enrichment = 1 denotes the null expectation. **(B)** Number of concordant DE gene–TE pairs within 5 kb, by TE class, timepoint, and strain. Concordance is defined as both the gene and TE showing the same direction of fold change within the same contrast × strain context. Only the four most frequent TE classes are shown; remaining classes are pooled as “Other”.
